# Lineage sorting inflates cross-environment functional enrichment

**DOI:** 10.64898/2026.08.04.742871

**Authors:** Kankan Zhao, Yiling Wang, Ran Xue

**Author notes:** Correspondence and requests for materials should be addressed to Kankan Zhao or Ran Xue.

## Abstract

“Everything is everywhere, but the environment selects.” What it selects has never been specified: genomes, lineages carrying their inherited gene content, or genes, rewritten inside lineages already in both places. Separating them needs a measurement, not a method: inflation is visible only where uncorrected and corrected estimates sit side by side on the same genes. Comparing only same-genus genomes across six catalogues, the median effect of the 178 orthologues a conventional soil–ocean analysis calls strongly enriched falls from 0.250 to 0.030, 45 reversing sign, DMSP demethylation keeping 2% of its reported effect and high-affinity phosphate transport reversing. On six host-niche contrasts within one *Escherichia coli* collection the same estimator changes almost nothing, retaining 92 to 100% with no reversals: the collapse belongs to the comparison, not the correction, its size varying over a hundredfold across eight contrasts. Fifty-nine orthologues survive family-wise permutation control, no negative control among them; they concern light, salinity, desiccation and metal efflux, and half are invisible to screening. Re-tested literature claims keep a median 24% of their effect but beat matched random sets 2.7-fold. Reading enrichment as adaptation requires lineage control and a per-contrast bound on what a comparison can manufacture from nothing.

## Main text

Baas Becking’s dictum, *everything is everywhere, but the environment selects*^1,2^, has been argued over for ninety years^3,4^, but what the environment selects has never been specified. The environment may select genomes, determining which lineages colonise and persist, with each carrying its entire inherited gene content into that environment; or it may select genes, rewriting gene content within lineages already established in both places. The two leave the same footprint in the comparison genome-resolved metagenomics has made routine: annotate a catalogue from each environment against a common scheme, compare carriage rates, and report that a gene is enriched in one. But a lineage that wins in one environment contributes more than a thousand genes to that environment’s enriched list, and for most of them the association reflects shared ancestry rather than function. Conventional enrichment analysis measures selection on genomes and reports it in the language of selection on genes.

We state the distinction operationally, because gene versus genome is otherwise a claim about units of selection that we do not make. What is separable in data is the level at which an association was established: between lineages, which is lineage sorting, or within them, which is within-lineage rewriting. Adaptation fixed deep in the tree is not separable from the first by any within-lineage design and we treat it as a residual.

The statistical problem is population stratification, which neighbouring fields addressed long ago: human association studies have corrected for ancestry for two decades^5^, and bacterial ones, facing a more acute form of it because clonal genomes inherit gene content in large linked blocks, condition on the clonal frame or the phylogeny^6–10^. Cross-environment functional enrichment, arguably the most common comparative use of genome catalogues, has largely not adopted the correction, and the winner’s curse compounds the resulting inflation because it is the top of an uncorrected ranking that gets followed up^11,12^. We deliberately use the estimator those fields standardised rather than a new one, so that the inflation we report cannot be attributed to a novel method. Population structure is the same confound in both settings; what differs is how much of it there is, and we measure that difference against the setting where the correction is already routine.

Related work has established the pieces: function and taxonomy decouple at community level^13,14^, trait conservation has a mapped phylogenetic depth^15,16^, variance has been partitioned between environment and phylogeny within species^17^, and enrichment has been tested for consistency across independent lineages^18,19^. What is missing is a measurement rather than a method. Inflation is observable only where the uncorrected and the corrected estimate sit side by side on the same features, and none of this work carries both arms: a partitioned variance ratio has no uncorrected arm to inflate, and of the two studies closest to our question one relates gene-content distances *between* genera to habitat annotations^20^, the between-lineage variation we attribute to sorting, while the other tests each niche within a species against the other niches rather than against an uncorrected baseline^21^. We ran the two-armed comparison on five global genome catalogues and one root isolate collection.

## Results

### Holding lineage fixed collapses cross-environment enrichment

We compared species-representative genomes from soil (SMAG^22^, 7,252) and ocean (GOMC^23^, 15,653) at completeness ≥ 70% and contamination ≤ 5%, annotated through one eggNOG-mapper pipeline, giving 8,077 KEGG orthologues present in at least 100 genomes. Effect size throughout is the between-environment difference in carriage rate, the proportion of genomes in which a feature is present rather than its abundance in reads or transcripts. Seventy-five genera had at least three genomes in each biome (1,682 genomes); within each genus, both the feature matrix and the biome indicator were residualised on annotated feature count, CheckM completeness and contamination (Methods).

Applied to the 178 orthologues a conventional analysis would report as strongly enriched (naive |Δ| ≥ 0.20), the design removed almost all of the signal. The median |Δ| fell from 0.250 to 0.030 (95% CI 0.021–0.038; Fig. 1b), a retention of 0.097 (0.069–0.133). Retention throughout is the median of the individual signed ratios, not the ratio of the two medians, which is 0.12; the two differ because a quarter of these features reverse sign. The collapse was not confined to the top of the ranking: across all 8,077 orthologues, regressing the genus-matched on the naive estimate gave a slope of 0.109 (0.083–0.130; Fig. 1a), so a conventionally reported difference of 30 percentage points corresponds within a genus to about three. Forty-five of the 178 reversed sign (37–61). Intervals throughout resample genera rather than genomes (Methods).

**Fig. 1.**
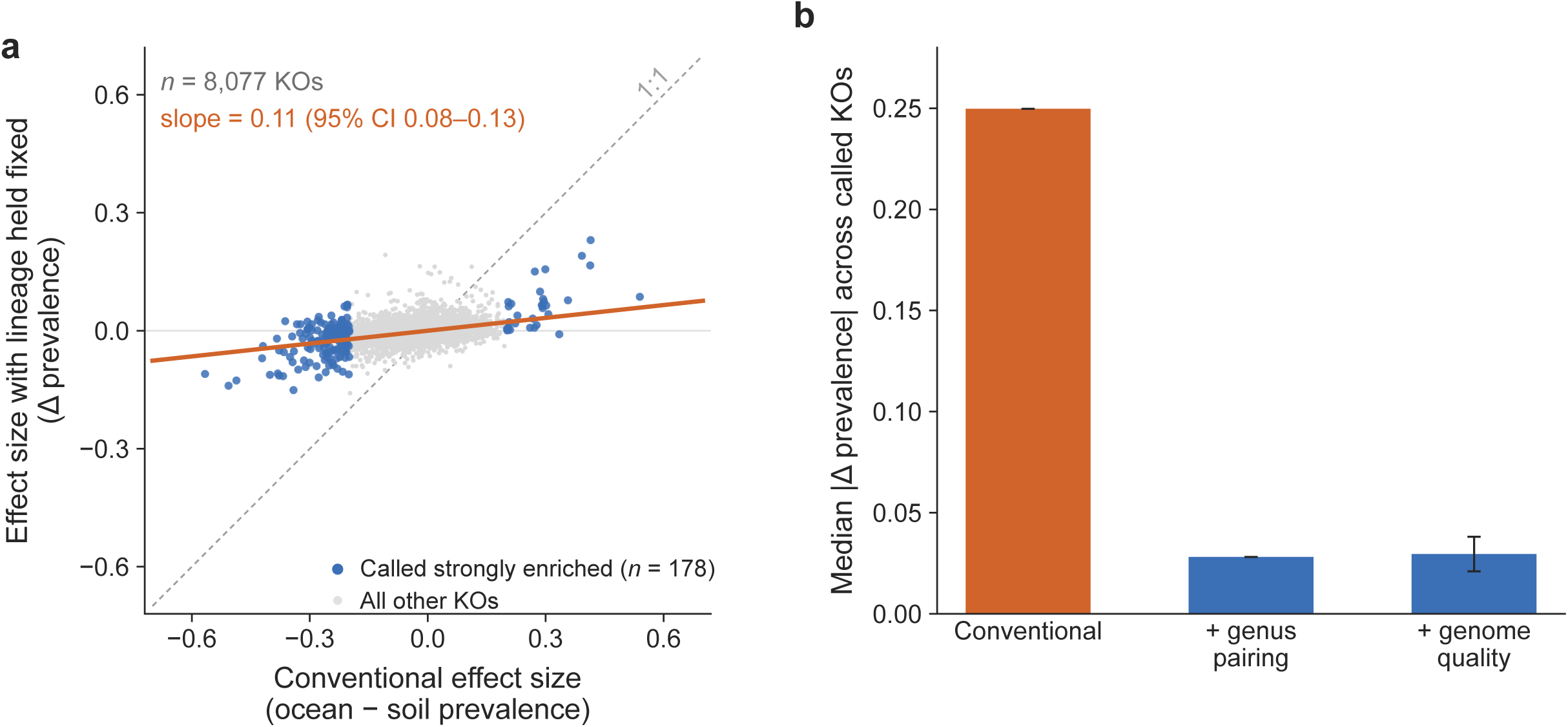
Holding lineage fixed collapses cross-environment enrichment. **a**, Naive versus within-genus controlled effect for all 8,077 KEGG orthologues in the soil–ocean comparison (*n* = 1,682 genomes, 75 genera). Each point is one KO; blue marks the 178 KOs a conventional analysis calls strongly enriched (naive |Δ| ≥ 0.20). The diagonal is no shrinkage; the fitted line is ordinary least squares through all 8,077 points, slope 0.109 (95% CI 0.083 to 0.130, 200 genus bootstraps). Points crossing the horizontal axis have reversed sign under control. **b**, Median |effect| for those 178 KOs at three levels of control: naive, within-genus pairing alone, and the main specification. The error bar is the symmetric normal interval, estimate ±1.96 bootstrap s.e. over 200 genus bootstraps (0.021 to 0.038). It is reported in this form rather than as a percentile interval because the statistic is a median of absolute values: resampling genera adds noise, |·| is inflated by noise, and the replicate distribution therefore sits above the point estimate, which falls at its 12.5th percentile. Replicate skew is mild (0.33). Every other bootstrap quantity in this paper is centred on its point estimate — the slope at the 52nd percentile of its replicates, retention at the 46th — and keeps the percentile interval. The first two bars carry no interval, because the naive quantity is fixed by construction and the pairing-only layer was not resampled.

An internal negative control shows why quality covariates are needed and defines a threshold applied to every subsequent result. Ribosomal proteins (92 orthologues) are universal and single-copy, so their true between-biome difference is zero by construction; genus matching alone left them at 0.020 and the controls brought them to 0.007, whereas without the controls the largest within-genus effects in the dataset were ribosomal (Extended Data Fig. 1a). What bounds the interpretation of any single feature is not this median but the upper tail. We therefore define the artefact floor of a contrast as the 95th percentile of |controlled effect| across genes whose true effect is zero, computed separately for each contrast; for soil–ocean it is 0.055 (95% CI 0.048–0.058). The residue that remains on the control, a mean *t* of +0.55, is assembly incompleteness rather than biology (*r* = −0.986 against completeness; Extended Data Fig. 1c).

The collapse is not lost power, genome size or pairing strictness (Extended Data Fig. 2), and it reproduced on 7,706 Pfam domains, where the median falls from 0.245 to 0.033.

### What survives family-wise control

We replaced the magnitude threshold conventionally applied to enrichment lists with a permutation test that has no free parameter, permuting the biome label within genus 10,000 times and recording the maximum standardised statistic across all 8,077 features to give a family-wise threshold of |*t*| = 4.458 (Monte-Carlo s.e. 0.008; Methods). Fifty-nine features exceeded it and none of the 92 ribosomal proteins did (Fig. 2a). All 59 also exceeded the artefact floor of their own carriage band, the two criteria bounding different errors and both being required (Methods).

**Fig. 2.**
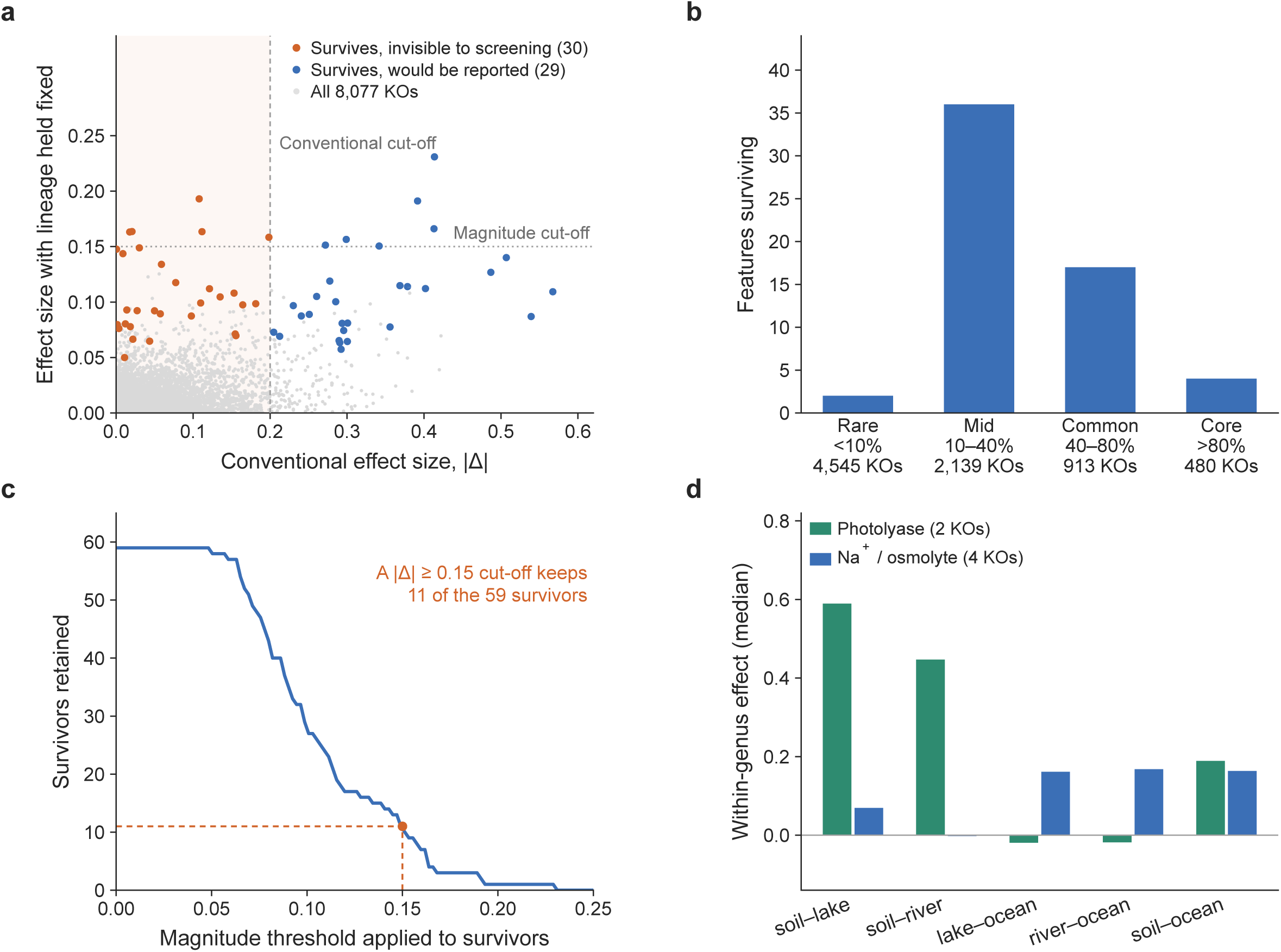
What survives family-wise control. **a**, The 59 surviving KOs positioned by naive and controlled effect, coloured by whether a conventional screen at naive |Δ| ≥ 0.20 would have reported them (29 would, 30 would not); grey is all 8,077 KOs. Survivorship is exceedance of a family-wise |*t*| threshold of 4.458, the 95th percentile of the maximum standardised statistic over 10,000 within-genus label permutations (Monte-Carlo s.e. 0.008). The shaded region and the dotted horizontal are the two cut-offs a conventional screen applies. **b**, Carriage-rate distribution of survivors against all tested features, in four prevalence bands; band totals are 4,545 rare, 2,139 mid, 913 common and 480 core features. **c**, Number of the 59 survivors retained as a function of a magnitude cut-off on |controlled effect|; at the |Δ| ≥ 0.15 threshold this literature commonly applies, 11 of 59 remain. **d**, Freshwater dissociates light from salinity: median controlled effects for the light axis (two DNA photolyases) and the salt and osmotic axis (four sodium and osmotic transporters) across soil to lake (7 paired genera, 120 genomes), soil to river (9, 179), lake to ocean (9, 173), river to ocean (32, 655) and soil to ocean. Lake and river are lit but fresh, ocean is lit and saline, soil is neither. Each bar is a median over the features of that axis; no test is performed across axes, the comparison being of sign and magnitude pattern rather than of a null hypothesis.

The count depends on the unit treated as exchangeable: between-biome patristic distances exceeded within-biome distances in 39 of 58 genera, and restricting permutation to phylogenetic sub-clades left the count unchanged at one cut-off and reduced it by 29% at a stricter one (Methods). Carried across to the full sample that puts the survivor count at 42 to 59.

Survivors were concentrated in the mid-frequency accessory genome, with 36 carried by 10–40% of genomes and 17 by 40–80%, against 4 in the core and 2 among 4,545 rare features (Fig. 2b). Their naive effects were small: 30 of the 59 (51%) had naive |Δ| < 0.20 and would not appear in a conventional enrichment list. Their controlled effects are smaller again, and only 11 of the 59 reach the |Δ| ≥ 0.15 that this literature commonly applies as a magnitude threshold, so such a threshold would discard 48 of them (Fig. 2c).

The set was largely, though not entirely, interpretable (Extended Data Table 2). On the ocean side the largest controlled effects were a sodium-coupled amino-acid transporter (alsT/agcS, +0.23), a putative transposase (+0.19), a DNA photolyase (phrB, +0.19), a cation:proton antiporter (yrbG, +0.17), a copper/silver efflux component (+0.16), a betaine transporter (betT, +0.16), sulfide:quinone oxidoreductase (sqr, +0.16), the second photolyase (phr, +0.16) and a sodium-dependent dicarboxylate transporter (SLC13, +0.15); on the soil side an uncharacterised protein (−0.16), a magnesium transporter (corA, −0.15) and the non-homologous end-joining protein Ku (−0.14). Six axes were represented: salinity and osmotic balance, light, desiccation, efflux of metals and metalloids (four genes, including a Zn² /Cd² -exporting ATPase and an arsenite transporter), defence (type I restriction, modification) and sulfur oxidation. Two of the largest effects, the transposase and the uncharacterised protein, belong to no axis.

Two survivors show what the lineage-controlled test adds rather than subtracts. A copper/silver efflux protein has a naive effect of 0.000 and a controlled effect of +0.148, and sqr moves from +0.018 to +0.163; the heavy-metal efflux claim as a whole is unauditable here precisely because its naive effect is −0.004, yet three of its genes survive. A conventional screen does not rank these low, it never sees them. To test whether each axis tracked its own physical driver rather than an overall marine–terrestrial gradient, we added lake^24^ and river^25^ catalogues, which are lit but not saline and were produced by independent groups (Fig. 2d). The photolyases had a median within-genus effect of +0.59 from soil to lake and +0.45 from soil to river, but −0.02 from either freshwater catalogue to the ocean; the four sodium and osmotic transporters showed the reverse pattern (+0.07 and −0.00 from soil to fresh water, +0.16 and +0.17 from fresh water to ocean); and Ku fell by 0.46 and 0.48 from soil to lake and river while remaining flat between the two aquatic biomes. The two freshwater catalogues gave near-identical values on both axes, while three candidates a conventional analysis would have promoted failed to reproduce (Fig. 2d).

### Re-testing the field’s standard adaptation claims

We assembled 27 adaptation claims of the standard form, spanning ion transport, osmolytes, light, nutrient limitation, metals, defence, degradation, motility and metabolism (Methods). Twenty-three were auditable in our data (naive |Δ| ≥ 0.02); all 27 are listed with sources and per-claim results in Extended Data Table 1.

The median retention ratio, controlled divided by naive effect, was 0.24 (quartiles 0.07 and 0.60; Fig. 3a,b). Nine of 23 claims kept their sign at *Q* < 0.05 and five reversed sign outright (Extended Data Table 1). Claims concerning salt and phosphorus limitation held up best (ectoine synthesis 1.56, phosphonate utilisation 0.71), while those concerning polymer degradation and specific marine metabolisms collapsed (chitinase 0.17, high-affinity phosphate transport −0.03, DMSP demethylation 0.02). One claim was specification-dependent: alkaline phosphatase retained 0.41 when only annotated feature count was controlled but reversed to −0.06 once completeness and contamination entered the residualisation, and we do not count it among the survivors.

**Fig. 3.**
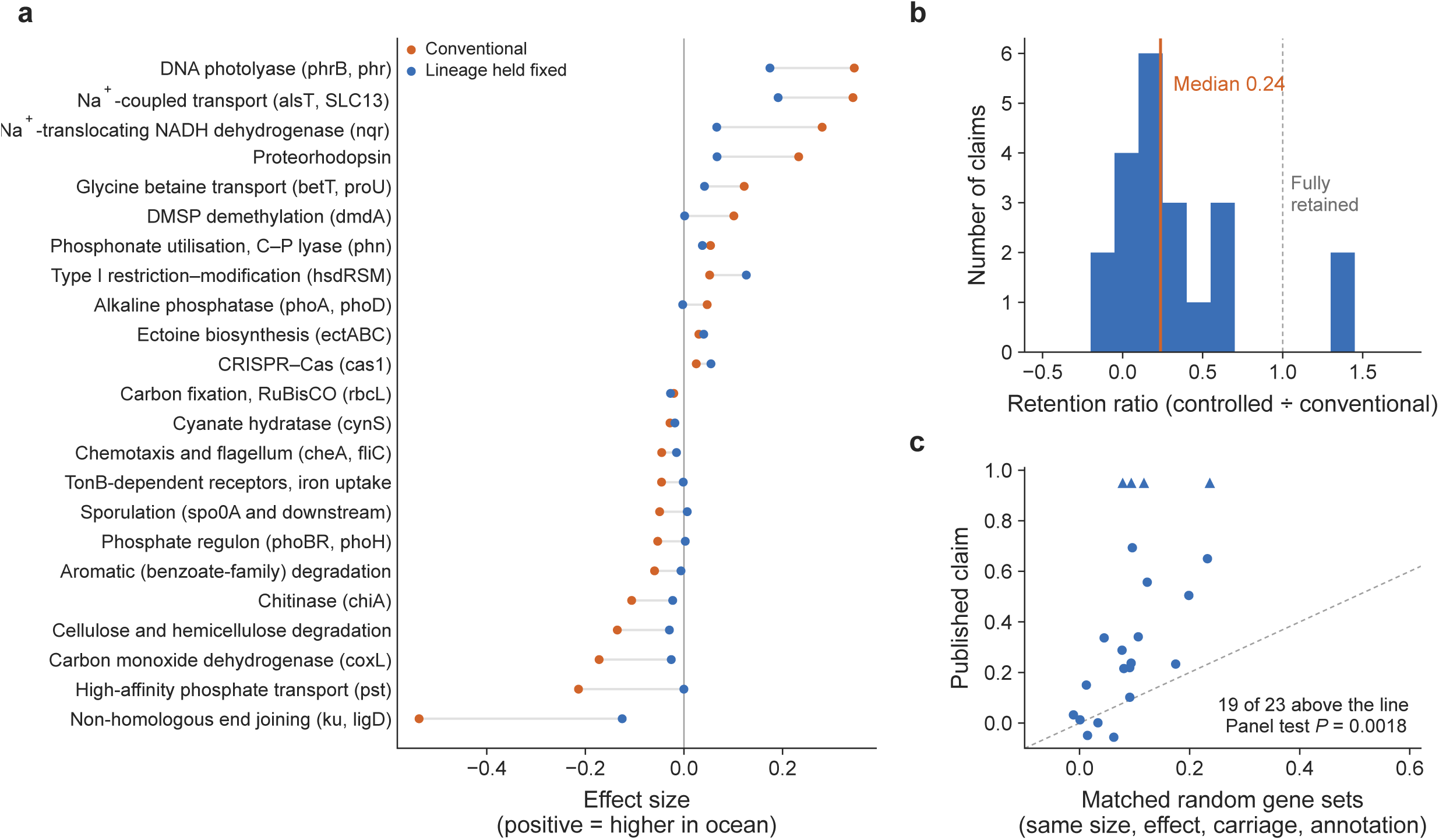
Re-testing the field’s standard claims. **a**, Naive and controlled effect for each of the 23 auditable claims of the 27 compiled, ordered by naive effect size. Claims whose two points fall on opposite sides of zero reverse sign under control (5 of 23). **b**, Distribution of the 23 retention ratios (controlled ÷ naive effect); the solid rule is the median, 0.24, and the dashed rule is a retention of 1, meaning no shrinkage. **c**, Observed retention of each claim against the median retention of its own matched null distribution, which matches claim size, naive effect, per-member carriage rate and annotation status. Nulls are 5,000 resamples. Four claims retain more than their naive estimate; they are drawn as triangles at the 0.95 ceiling of the *y* axis rather than at their own values, which would compress the rest. The reported *P* value, 0.0018, is a panel-level permutation *P*, the fraction of 5,000 matched null panels whose median retention reaches the observed panel median; it is one test on the panel, not 23 independent tests. Claim-panel effects are the three-covariate main specification, the same standard as the rest of the paper; the feature-count-only values, which several panel members being Pfam features argue for, are in Extended Data Table 1.

A retention of 0.24 is consistent both with claims that are largely artefactual and with claims that are real but miscalibrated, and the two differ in what a random gene set would have achieved. Against null sets matched on the four properties that trivially move a retention ratio, claim size, naive effect, per-member carriage and annotation status (Methods), the 23 claims retained 0.237 where matched random sets retained 0.089, that figure being the median across resampled panels rather than the median of the per-claim nulls listed in Extended Data Table 1, which is 0.091. Because the claims are not independent draws the test was run at panel level (*P* = 0.0018; Fig. 3c), and nested subsets at naive |Δ| ≥ 0.10 and ≥ 0.20 gave *P* = 0.0028 and 0.0014. On the ratio-free statistic, the controlled effect itself, claims reached a median 0.027 against 0.011 (*P* = 0.0002). Dropping each claim in turn left a worst-case *P* of 0.0028. Retention is quoted at three values in this paper, 0.10, 0.24 and 0.29, and they are one statistic on three populations rather than three results: features a conventional analysis calls strongly enriched, published claims, and the variant sets an association study’s own filter passes. A retention ratio rises as the population it is computed over is selected less extremely, so the three are ordered as selection on the naive effect weakens. The claims do not all rest on the same kind of original evidence, and a claim made on read abundance is not the quantity we re-test; restricted to the six whose original evidence was genome prevalence, the same quantity, retention is 0.18 against 0.24 for the whole panel, close enough that the shrinkage is not an artefact of quantity mismatch (Extended Data Table 1).

The correction does not remove true signal: as a positive control, the sodium-translocating NADH dehydrogenase nqr, supported by ancestral-state reconstruction in an independent set of 11,248 aquatic MAGs^46^, retained sign and significance (+0.280 to +0.068, *P* = 0.0004) at 24% of its reported magnitude. Pathway-level analysis fared worst: of 239 KEGG modules, 34 reached |Δ| ≥ 0.05 conventionally and none survived. This follows from the composition of the surviving set, since individual transporters do not constitute a pathway.

The surviving features are also where a classifier’s information lies. An L2-regularised logistic classifier predicting biome from gene content, with entire genera held out across eight folds, reached AUC 0.819 on KEGG features and 0.853 on Pfam against 0.563 and 0.543 for within-genus permuted labels (Fig. 4). Deleting the features the per-feature test selects, with every deletion paired against a matched random deletion and the selection refitted inside each fold so that it never sees the genomes it is scored on, cost 0.080 of AUC out of sample (0.740 against a matched-random 0.820 ± 0.003); fitted on all genera instead the same deletion cost 0.234, so two-thirds of the apparent effect was selection reaching the test fold, and Pfam agreed at both levels of rigour (Methods). That out-of-sample cost is a third of the classifier’s entire 0.256 margin over permuted labels, against a matched random deletion that costs nothing. Deleting only the 59 family-wise survivors, a set twenty times smaller, costs 0.017 out of sample on KEGG features and 0.024 on Pfam, so the surviving set is a concentrated but small part of the recoverable signal. The decline with rank is smooth rather than stepped (Fig. 4b): no small group of features carries the signal alone, and forward selection saturates at roughly 100 features only because the top features are mutually redundant (median maximum correlation 0.50). None of this reinstates conventional screening, since half the features the lineage-controlled test selects have naive effects too small to appear in a conventional enrichment list.

**Fig. 4.**
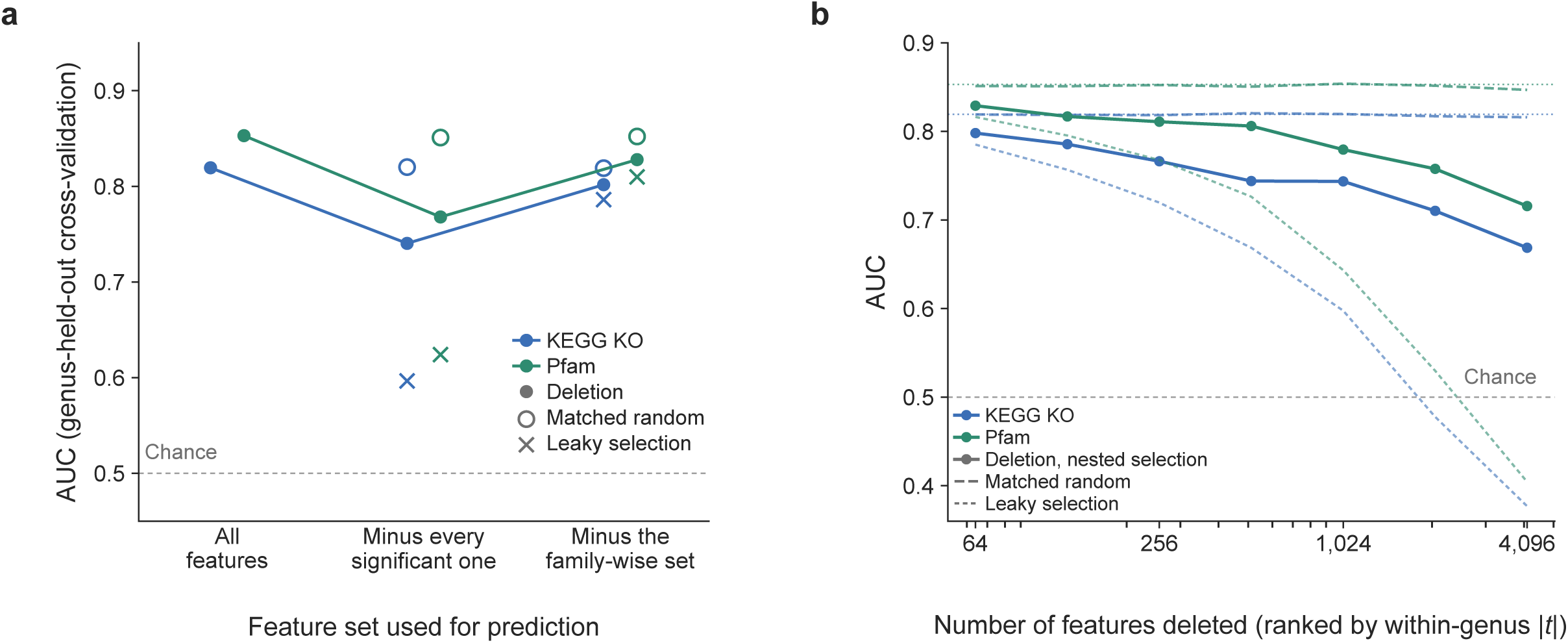
Where the classifier’s information lives. L2-regularised logistic regression (C = 0.05) predicting biome from binary feature vectors, with entire genera held out across eight folds, so no genus appears in both training and test; all AUCs are pooled over held-out predictions. **a**, AUC for the full feature set, for the per-feature significant set deleted (*q* < 0.05, mean 984 features across folds), and for the family-wise survivors deleted (mean 48 features). Filled circles joined by a line are the nested analysis, in which the deletion set is re-selected inside each fold from that fold’s training genera alone. Open circles are the matched random deletion, the same number of features drawn to match the deleted set’s carriage distribution over 20 prevalence bins, mean of 20 draws (s.d. 0.003 or less). Crosses are the same deletion with selection fitted on all genera; the gap between cross and filled circle is selection leakage. AUC is drawn on a truncated axis with the chance line at 0.5 marked, and as points rather than bars because AUC has no meaningful zero against which bar heights could be read as ratios. **b**, AUC after deleting the top *k* features ranked by within-genus |*t*|, for *k* = 64 to 4,096, under fold-wise ranking (solid), all-genera ranking (fine dashed) and matched random deletion (dashed). Dotted horizontals mark the full-feature AUC. Beyond *k* = 1,024 more than an eighth of the matrix is gone and the curve is not read as a statement about a feature set.

### The scope of the result across contrasts and catalogues

The collapse appeared at every analytical level we could measure and is therefore not a property of the annotation layer (Fig. 5a). Median retention was 0.10 across the 178 orthologues called strongly enriched, 0.24 across the literature claims and 0.06 across 239 KEGG modules. Whole-proteome traits computed from sequence rather than annotation, which share no methodology with the gene-content analysis, collapse furthest of all: a median retention of 0.02 across nine traits, the largest being nitrogen atoms per residue at 0.27 (Extended Data Fig. 3). Retention therefore varies about tenfold across levels rather than being constant, and in an interpretable direction, since the claim level is the only one whose members were chosen by someone for being biologically important.

**Fig. 5.**
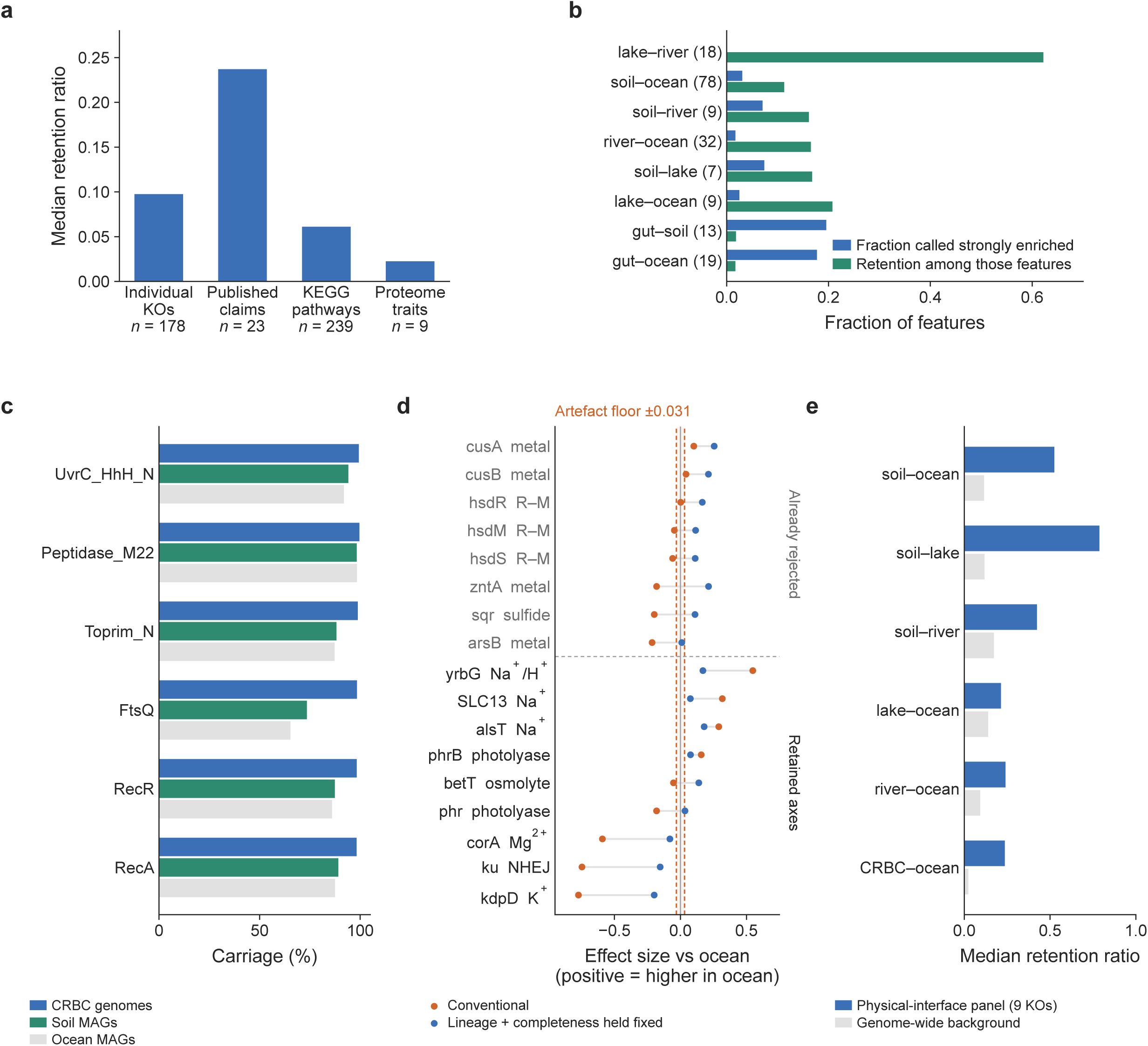
The scope of the result across contrasts and catalogues. **a**, Median retention at four analytical levels: the 178 individual KOs a conventional analysis calls strongly enriched, the 23 auditable published claims, 239 KEGG modules and 9 whole-proteome traits. Every level collapses; the sequence-based traits, which share no methodology with the gene-content analysis, collapse furthest. The module level is not an independent measurement, being by construction the mean of its member genes’ effects (Methods). No test is shown, these being four descriptive medians on different quantities. **b**, For all eight biome contrasts, the fraction of Pfam features a conventional analysis calls strongly enriched (naive |Δ| ≥ 0.20) and the median retention among those features, ordered by genus-level Jaccard overlap between the two catalogues; paired genus counts are given on the axis. Lake against river, two freshwater catalogues built by independent groups, is the design-level negative control at the high-overlap end; the human gut is the extreme at the other. The eight contrasts are not mutually independent, two sharing the gut catalogue and four sharing soil or ocean, which is why the overlap relationship is reported without an interval (Extended Data Fig. 4). **c**, Carriage of six universally present genes in the rhizosphere isolate collection against the two MAG catalogues, the sanity check run before any comparison; percentages are carriage across each catalogue’s paired genomes. **d**, Naive and controlled effects for the nine physical-interface orthologues in the CRBC, ocean contrast (71 paired genera, 3,621 genomes). The dashed verticals are that contrast’s artefact floor, the 95th percentile of |controlled effect| across the ribosomal and RNA polymerase orthologues whose true effect is zero, at 0.031 (95% CI 0.019 to 0.037 from 4,000 resamples of that gene set); features must clear the interval’s upper bound, not only the point estimate. **e**, Panel-level retention against the genome-wide background across all six CRBC contrasts. Panel-level rather than genome-wide retention is compared because the catalogues differ in compositional distance (Extended Data Fig. 6a).

It did depend on which environments were compared (Fig. 5b). On Pfam features and a common pipeline, retention ran from 0.11 for soil–ocean to 0.21 for lake–ocean, and fell to 0.02 against the human gut. The quantity that varied most was not retention but the size of the problem retention corrects: the fraction of features a conventional analysis calls strongly enriched ran from 0.08% to 20% across the same contrasts.

Soil–ocean, at 3%, is a point on that range rather than a representative value, and is our main contrast only because it is the sole pair with enough shared genera to support a maximum-statistic test. The range was ordered by taxonomic overlap: genus-level Jaccard overlap was negatively correlated with the fraction of features called enriched (rho = −0.71, *P* = 0.047; Extended Data Fig. 4), whereas the correlation with retention itself was not significant (rho = +0.52, *P* = 0.18), so we state the dose–response only for the quantity that carries it. Eight non-independent contrasts make this a rule of thumb rather than a calibration curve.

The two ends of the range each supply a control the design otherwise lacks. At the low-turnover end, lake against river is a negative control at the level of the design rather than the feature: both catalogues are freshwater, so no environmental contrast separates them, yet they were built by independent groups, so any apparent enrichment between them is catalogue batch effect and lineage composition alone. Their genus overlap was the highest of any pair we ran (Jaccard 0.157, 18 paired genera) and a conventional analysis called 0.08% of features strongly enriched. Retention there is 0.62, the highest of the eight contrasts, which is what the quantity should do where there is almost nothing to correct: the ratio is computed over the handful of features that clear the conventional threshold at all, and it is the numerator, not the ratio, that carries the comparison. Had the collapse we report been an artefact of comparing two independently built catalogues, it would have appeared here. At the high-turnover end the human gut is the extreme case (Extended Data Fig. 5), with only 13–19 shared genera, 18–20% of features called strongly enriched, and a retention of 0.02 that both known biases there make an upper bound.

Every result so far rests on MAGs, in which incompleteness is controlled statistically rather than eliminated. The Crop Root Bacterial Genome Collection^27^ (CRBC) is a partial escape from that confound, with 3,370 of its 4,391 paired genomes isolate assemblies and mean completeness 96.1%, though one in five falls below 95% (Fig. 5c, Methods). Read against this contrast’s own artefact floor of 0.031 (95% CI 0.019–0.037), all nine physical-interface orthologues cleared both the floor and its upper bound (Fig. 5d; naive to controlled: kdpD −0.77 to −0.21, alsT +0.29 to +0.19, yrbG +0.55 to +0.17, Ku −0.75 to −0.16, betT −0.05 to +0.14, corA −0.59 to −0.09, phrB +0.16 to +0.09, SLC13 +0.32 to +0.08, phr −0.18 to +0.04), though two did so only after reversing the sign of their naive estimate here, betT and phr, and we would not report those on this contrast alone. The control is the comparison without a biome contrast in it: CRBC and soil are both terrestrial, and against that contrast’s floor of 0.094 only two of the nine cleared the line. Against its own background the panel exceeded genome-wide retention in all six contrasts by 1.5- to 10.1-fold (Fig. 5e), most widely where assembly quality is highest, which is the opposite of what incompleteness would produce.

### How far down the tree the result holds

Genus is one threshold on a continuum of relatedness, and residual within-genus structure could still inflate the estimates. We built a maximum-likelihood tree from 25 concatenated ribosomal proteins (1,153 genomes, 56 testable genera) and added a phylogenetic random effect on the Gower-centred patristic kinship matrix under the main specification (Fig. 6a). Biome clustered phylogenetically within genera, but weakly (median relative excess +3.7%). The ladder was 59, then 28, then 25: restricting to genomes on the tree cost 31 features and adding the phylogenetic random effect cost 3 more, so the loss is almost entirely sample size and phylogeny below the genus is nearly free. The shared phylogenetic variance ratio is not identified, so the survivor count moves between 18 and 28 across the range of h² the data cannot distinguish (Fig. 6b). The reason bears on interpretation: per-feature h² is bimodal, with single-copy core genes at 0.97 and mobile elements at 0.003 (Fig. 6c), and the 59 survivors sit at the vertical end (median h² 0.909, indistinguishable from carriage-matched background, *P* = 0.09). The survivor set is itself bimodal, however: two of its sixteen largest effects, the putative transposase and the type I restriction, modification S subunit, sit at h² = 0.003, and two copper/silver efflux components at 0.17 and 0.44.

**Fig. 6.**
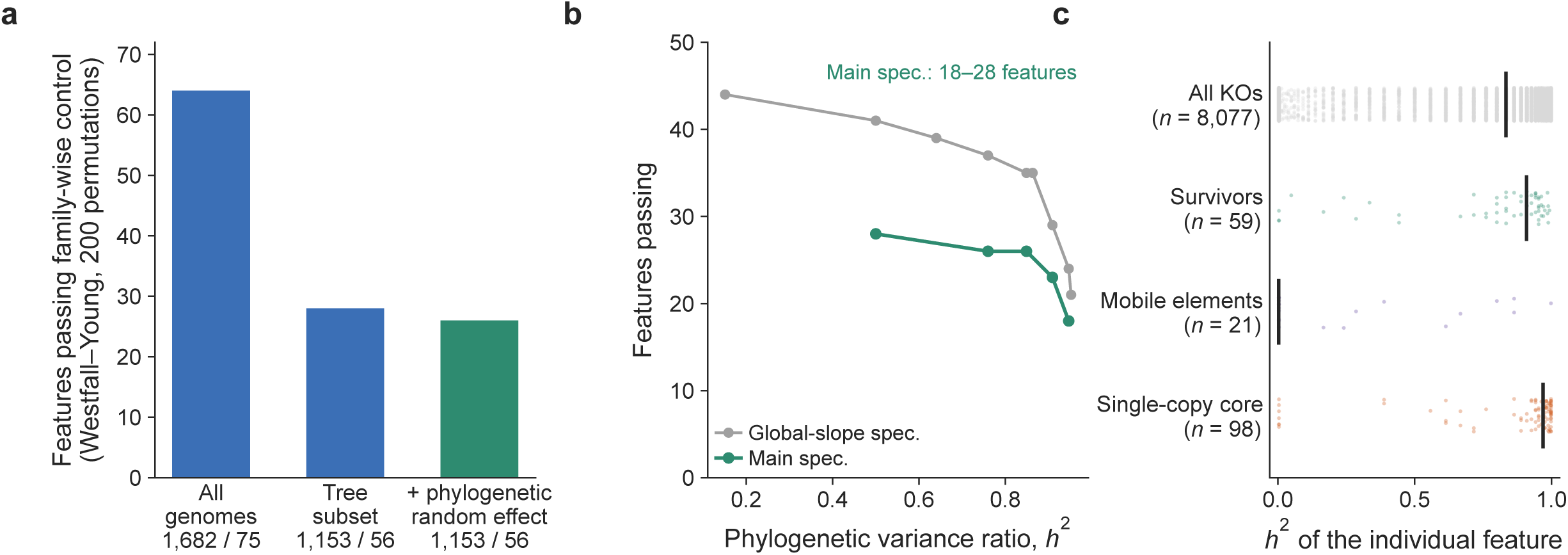
Tightening lineage control below the genus. **a**, Number of features passing the family-wise threshold at each rung: genus-matched on the full sample (1,682 genomes, 75 genera), restricted to the genomes on the ribosomal tree (1,153 genomes, 56 genera), and with a phylogenetic random effect added on the same subset. The threshold is re-estimated by 10,000 permutations at each rung, so the rungs are comparable. Separating the two steps separates the cost of sample size from the cost of phylogeny. **b**, Features passing across the grid of the shared phylogenetic variance ratio h², which the data cannot identify, under the main specification and under a global-slope specification; the main specification spans 18 to 28. **c**, Per-feature h² at its REML optimum for single-copy core genes, mobile elements, the 59 survivors and all KOs; vertical rules are group medians and *n* is given per group. The survivor median, 0.909, is indistinguishable from a carriage-matched background (*P* = 0.09), but the survivor distribution is itself bimodal.

Effects persisting within a genus but not within a species would read as sub-clade sorting rather than as gene content rewritten in place, but that design cannot be run here: only 73 named species occur in both catalogues and none has three genomes on both sides, because each catalogue is dereplicated to one representative per species. A paired comparison of 74 species pairs is possible and is uninformative in both magnitude and direction.

Both failures trace to one bias the design cannot absorb. The two sides of a pair differ by 3.7 percentage points of completeness, and a stratum of two genomes cannot support a within-stratum quality adjustment; adjusting across pairs moves the mean ribosomal *t* from +1.24 to +1.42 rather than towards zero. The artefact floor is consequently 0.219 against a median within-species effect of 0.0135, sixteen-fold above the signal, so no magnitude is supportable, and power is not the constraint (at a median standard error of 0.013, a genus-level effect of 0.098 carried intact would have been detected). Sign preservation is not interpretable either, because the completeness gap is directional and the apparent effect sits entirely in the ocean-positive half it would inflate, disappearing when pairs are matched on completeness (Extended Data Fig. 8).

Our catalogues cannot support a within-species design, but other people’s data can, and this is where the size of the problem separates from the machinery used to measure it. Applying the same estimator to the study closest to ours^21^, six host-niche contrasts within one *Escherichia coli* collection, holding lineage fixed retains between 92% and 100% of the uncorrected effect in every one of the six and reverses no signs at all. On the identical statistic and the same genomes, soil against ocean retains 29% and reverses 28%. The gap survives matching the two settings on lineage imbalance, where their contrasts and ours overlap, and on the number of genomes per lineage stratum, which we vary from 39 down to 16 against our 22 (Methods). The collapse is therefore not something this estimator produces wherever it is applied. It is a property of the comparison. Their unit is a k-mer and ours is an orthologue, so the two are matched on estimator, scale and stratification but not on the feature itself, which is a caveat on the comparison and not on its direction.

## Discussion

The environment selects genomes more than genes: at every level we can measure, most of what a conventional cross-environment comparison reports as functional enrichment is an accounting of which lineages live where. Two shrinkage factors run through this and should not be conflated. For an individual gene, retention is 0.097 (95% CI 0.069–0.133), so about nine-tenths of a reported effect is lineage accounting; for a published claim, a composite of genes chosen because someone thought it mattered, retention is 0.24, an overstatement of roughly fourfold. The gap is itself a result: claims are composites of widely carried, well-characterised genes, and both properties raise retention. Nor is the problem’s size a constant: the fraction of features called enriched varies by more than two orders of magnitude across our contrasts, ordered by a taxonomic overlap that can be computed before any gene is tested, though on eight contrasts that share catalogues this orders the comparisons rather than predicting a value for any one of them. Carriage is moreover the conservative case, since an abundance-weighted comparison lets one dominant lineage contribute its whole gene complement in proportion to its abundance, making the hitchhiking fraction there larger than the one we measure.

We do not conclude that the literature’s findings are false. Effect sizes are overstated, a minority reverse sign, and the pathway, the level at which the field most often reports results, is the level at which nothing survives. But the audited claims exceeded matched gene sets by 2.7-fold in retention and 2.4-fold in controlled effect, so the field has been identifying real signal and mis-attributing its cause. A recent precedent indicates how such an audit lands: confounder control removed *Fusobacterium nucleatum* from the colorectal cancer microbiome while six other species held^47^. Partial collapse with partial survival is the expected outcome, not a sign of over-correction. What fails is the ranking rather than the testing. The effects environment leaves within lineages are individually small (median controlled effect 0.066 among the top hundred classifier features) but not undetectable; what loses information is the magnitude cut-off applied on top, which discards 48 of our 59 survivors. Pathway analysis fails for the same arithmetic reason, since a module-level statistic averages one or two real signals against dozens of hitchhikers.

The sharpest reading of our own result is one it does not support. The bulk of the surviving gene content is as phylogenetically structured as any comparable gene (median h² 0.909), and a feature organised that strongly by phylogeny is not being gained and lost repeatedly within a genus, so most of the layer resembles sorting among sub-clades more than gene content rewritten in place. But the two mechanisms coexist inside the surviving set rather than competing for it: its h² distribution is bimodal like the genome’s, and two of its largest effects, a putative transposase and a restriction, modification subunit, sit at the mobile-element end at h² = 0.003. Genome-level and gene-level selection are therefore not a dichotomy with a natural boundary but a continuum, on which we have measured one cut. The reading is scale-dependent: within one species, host-associated rewriting is carried predominantly by mobile elements^21^, whereas across genera and biomes what remains is vertically structured, as expected if plasmid-borne rewriting is lost on the longer timescale or fixed with its sub-clade. Below the genus the answer stops for a stateable reason: a phylogenetic random effect within genera costs a tenth of the surviving set, one rank further is impossible on catalogues dereplicated to one genome per species, and on the species pairs that do exist the artefact floor rises sixteen-fold above the signal. That is a property of the catalogues rather than of the question.

Why the survivors are interfaces with the physical world rather than metabolism is a hypothesis we mark as one: light, salt, water and phosphorus limitation are stresses a cell can neither avoid nor delegate, whereas metabolic capabilities can be supplied by other community members. Four limitations bound the conclusions. Genera shared across environments are cosmopolitan and likely generalist, a bias running towards finding within-lineage adaptation and so strengthening a negative result; adaptations fixed deep in the tree are invisible to a within-genus design; every feature is presence/absence, leaving copy number, expression and regulation outside the design, so for claims originally established on metagenomic abundance we test whether a gene is carried rather than whether it is abundant; and assembly quality is controlled but not eliminated, so no MAG-based comparison can claim the confound is gone.

Reading cross-environment enrichment as adaptation therefore requires three things that cost no new data: lineage control, an artefact floor computed per contrast rather than one global threshold, and a permutation-based family-wise threshold in place of a magnitude cut-off. Supplementary Table 1 sets these out with the rest of what our own audit turned up, each item against the evidence for it. Whether the within-lineage layer shrinks below the genus, and whether the concentration of surviving adaptation at physical interfaces reflects the delegability of metabolic function, are the two questions that do require new data.

## Methods

### Genome catalogues

Six catalogues were used, each downloaded as the authors published it and re-processed here from protein sequence onward. Soil: SMAG^22^, 21,077 species-level genome bins from 3,304 soil metagenomes, of which 7,252 met our quality filter and carried a genus assignment. Ocean: GOMC^23^, 24,195 species-level genomes from marine metagenomes collected 2009–2020, of which 15,653 passed. Lake: the LakePulse catalogue^24^ of 1,008 genomospecies from 308 Canadian lakes, 293 passing. River: GROWdb^25^, 1,469 species from 163 samples across 106 North American river sites, 920 passing. Human gut: UHGG species representatives^26^, 4,326 passing. Rhizosphere: the Crop Root Bacterial Genome Collection (CRBC)^27^, 6,699 genomes from wheat, rice, maize and *Medicago* roots.

### Quality filter and its limits

Genomes were retained at CheckM^32^ completeness ≥ 70% and contamination ≤ 5%, the MIMAG^35^ medium-quality boundary. This threshold is permissive by design: raising it removes genomes non-randomly with respect to biome and lineage, so the confound is handled statistically in the main analysis and by subsetting as a sensitivity check rather than by a strict filter (see *Sensitivity to assembly quality*). Contamination values were taken from the source catalogues’ own CheckM output (Extended Data Table 2 of ref. 22 and Supplementary Table 1 of ref. 23, not the tables of that name in this paper) rather than recomputed, so they inherit those pipelines’ CheckM versions.

We state one correction to how the CRBC collection has been described, including by us in an earlier draft. It is not a set of complete isolate genomes. Of the 4,391 CRBC genomes entering our paired analysis, 3,370 are isolate assemblies and 1,021 are MAGs; mean completeness is 96.1% (median 99.5%, minimum 50.3%, 20% below 95%), the median contig count is 30, and every genome is classified MIMAG medium quality. CRBC is therefore a higher-quality catalogue than the MAG sets, not a confound-free one, and we report it that way throughout.

### Taxonomy and the definition of a lineage

Genus labels are GTDB^33^ labels as assigned by each source catalogue; GOMC is annotated against GTDB r207 and SMAG against the release current at its publication. Lake genomes were classified locally with GTDB-Tk v2.4.1^34^ against the r226 reference package. Because GTDB genus boundaries are set by relative evolutionary divergence rather than by nomenclature, a genus is a comparable unit of relatedness across catalogues, which is what the design requires; it is not a claim that genus is the biologically correct scale, and the phylogenetic mixed model and the within-species analysis below test scales above and below it. Genomes without a genus assignment cannot enter a within-genus design and were dropped. Catalogues are independently dereplicated to species representatives, so within one catalogue a species contributes at most one genome, a fact that constrains the within-species analysis below.

### Uniform functional annotation

All protein sets were annotated with a single eggNOG-mapper^28^ installation (emapper-2.1.12, DIAMOND 2.1.8) against emapperdb-5.0.2^29^, at default sensitivity and E-value, with taxonomic scope set to auto. Published annotations were never mixed in: UHGG and CRBC were re-annotated from protein sequence with the same installation. Features are KEGG^30^ orthologues and Pfam^31^ domains, scored as presence/absence per genome. A KO or domain enters the analysis if it is present in ≥ 100 genomes of the contrast (≥ 50 for the smaller CRBC contrasts). Never comparing published annotations directly is not a formality: using UHGG’s native annotations, RecA, universal by definition, appears in 0.0% of gut genomes, because Pfam had renamed the domain RecA to RecA_N. Matching by accession recovers 96.3% and re-annotation through our shared pipeline gives 95.8%, while ribosomal domains match under either scheme, so the failure is both silent and selective (Extended Data Fig. 5a). This carriage filter is the reason some textbook marine adaptations are absent from our tables: proteorhodopsin KOs K04641 and K04643 are carried by 16 and 47 ocean genomes respectively out of 22,905, far below the threshold, so they are never tested rather than tested and rejected.

### KEGG modules

No presence threshold is applied, and none is needed. A module is scored per genome as the *fraction* of its constituent KOs that are present, a continuous value in [0, 1], and the module-level effect is the between-biome difference in that fraction, estimated by the same within-genus residualisation. Because the estimator is linear, this fraction-based module effect is exactly the mean of its member KOs’ effects, so the module-level result is a deterministic consequence of the gene-level one rather than a separate measurement with its own tuning parameter. Readers should note that this is the reason no module survives: averaging one or two real signals against dozens of hitchhikers is an arithmetic operation, not a statistical failure. The carriage filter is relaxed to ≥ 30 genomes for the module analysis so that sparsely carried module members are not silently dropped from their modules.

### Genus-matched design

Genera with ≥ 3 genomes in each environment were retained. The feature matrix and the biome indicator were both residualised, within genus, on the total number of annotated features per genome, CheckM completeness and CheckM contamination (Frisch–Waugh–Lovell^41,42^); the coefficient of the residualised indicator is the controlled effect. Residualising within genus gives each genus its own covariate slopes, which matters: a specification with slopes pooled across genera carries twice the residual bias on the ribosomal control (mean t +1.03 versus +0.55). Genera too small to support the covariate fit are centred only, which leaves the covariate uncontrolled rather than fitting it to noise. Per-feature *q* values are Benjamini–Hochberg^40^ adjusted. Standard errors are the ordinary least-squares errors of the residualised regression with degrees of freedom reduced by the genus-by-covariate parameters absorbed; they are not additionally clustered by genus, because the within-genus centring has already removed the genus-level random intercept that clustering would otherwise absorb. Uncertainty that a cluster-robust error would capture is instead reported by the genus bootstrap below. Implemented in lineagectl, which is unit-tested against simulated catalogues with known ground truth.

### Survivor criterion

Family-wise threshold by Westfall–Young^39^: the biome label is permuted within genus 10,000 times, the maximum |t| across all features is recorded, and the 95th percentile of that null is the threshold. Standardised statistics are used because the maximum of raw effect sizes is dominated by mid-prevalence features. Permutation is performed under the same design that produced the estimates. The Monte-Carlo error of the threshold is reported alongside it, because the survivor count is a step function of the threshold and an under-resampled quantile propagates directly into that count: at 200 permutations, which is what an earlier version of this analysis used, resampling the null moves the threshold by enough to move the survivor count by roughly ±10.

The permutation’s exchangeability assumption deserves a caveat we can quantify. Permuting the biome label freely within a genus assumes genomes of that genus are exchangeable with respect to biome, and they are not entirely: between-biome patristic distances exceed within-biome distances in 39 of 58 genera, significantly so in 16. This makes the within-genus permutation mildly anti-conservative, so the true threshold is somewhat higher and the survivor count somewhat lower than reported. We bound the size of the effect directly. Holding the estimator fixed, each genus on the ribosomal tree is cut into sub-clades by average-linkage clustering of its internal patristic distances at a fixed fraction *f* of the genus’s maximum internal distance, and the label is permuted only within sub-clades; *f* = 1 recovers free within-genus permutation. At *f* = 0.75 the survivor count is unchanged, and at *f* = 0.5 it falls by 29%. At *f* = 0.25, 87% of blocks hold a single biome and the permutation retains almost no freedom, which inflates the threshold to 7.42 and is reported as a degenerate design rather than as a corrected estimate. We therefore quote the survivor count as an upper end with a range extending about 30% below it, 42 to 59. The phylogenetic mixed model below is the second, independent check on the same question.

### Confidence intervals

Intervals are non-parametric bootstrap percentile intervals from 200 resamples in which *genera*, not genomes, are the resampling unit; each drawn genus is relabelled so that a genus sampled twice contributes two independent strata, which is the correct analogue of per-genus covariate slopes under resampling. Genomes within a genus are not exchangeable with each other, so a genome-level bootstrap would understate the interval^44^. The naive contrast is a whole-catalogue quantity and is held fixed across resamples; only the within-genus layer is resampled. The bootstrap estimator is checked against the main fit on the unresampled data and must reproduce it to within 10 before the resamples are run.

### Internal negative control and the artefact floor

Ribosomal protein KOs (K02863–K02999), universal and single-copy, have a true between-biome difference of zero by construction. We report their controlled effect as a median, an upper tail and a mean t. The median answers “is the estimator unbiased on average”; the upper tail answers the question that actually matters for interpreting any single feature, namely how large an effect this comparison can manufacture from nothing. We therefore define the artefact floor of a contrast as the 95th percentile of |controlled effect| across the union of two gene sets whose true effect is zero, the 92 ribosomal KOs and the RNA polymerase subunits, and require that a reported feature exceed its own contrast’s floor. The floor is contrast-specific and varies by threefold across our comparisons, so a magnitude that is interpretable in one contrast is not in another. Because it is a 95th percentile of roughly a hundred genes, it is reported with a bootstrap confidence interval obtained by resampling that gene set 4,000 times; relative interval widths run from 15% to 60% across our contrasts, the widest being the CRBC–ocean floor, and features are required to clear the interval’s upper bound and not only the point estimate. The floor and the Westfall–Young threshold are complementary rather than alternative: the first bounds systematic bias on the effect-size scale, the second bounds multiplicity on the *t* scale.

Because the floor is calibrated on genes whose prevalence is near one, where the sampling null is narrowest, it might be expected to be too lenient for the mid-prevalence features that dominate the surviving set. We tested that directly rather than assuming it either way, by computing the floor a second way: permuting the biome label within genus 2,000 times and taking the 95th percentile of |controlled effect| within each of ten bins of pooled carriage rate, so that every feature is judged against a null of the right width for its own prevalence. The permutation floor does peak at mid prevalence, as the sampling argument predicts, but it peaks at 0.032, against 0.055 for the single-copy set, so on sampling alone the scalar is the stricter of the two. The two quantities are not the same thing: a permutation null measures sampling noise at a given prevalence, whereas the single-copy set carries sampling noise plus the assembly-incompleteness bias diagnosed above.

The single-copy set also has a limit that should be stated rather than assumed away. Its members are distributed over carriage as their annotation recovers them, and none of the 92 falls between 10% and 40% carriage, which is where 36 of the 59 survivors sit; in that band the scalar cannot be calibrated at all and only the permutation null is available. Where the two can be compared the scalar is slightly the more lenient: restricted to the 27 single-copy genes carried by 40 to 80% of genomes, the band’s own 95th percentile is 0.059 against the scalar’s 0.055, and the magnitude of what a null gene reaches does scale with its carriage (Pearson *r* = 0.56 between carriage and |controlled effect| across the 92). We therefore tested every survivor against the strictest floor its own band admits, the larger of that band’s single-copy percentile and its permutation threshold. All 59 clear it, including all 17 in the 40 to 80% band against 0.059 (Extended Data Fig. 1d). We report both for every contrast where both can be computed. The dependence of the mean t on assembly completeness was measured by drawing subsets that hold the number of genomes, the genus composition and the two-sided balance of each genus exactly fixed while selecting the most complete, random, or least complete genomes within each cell.

### Sensitivity to assembly quality, and what the residual bias on the negative control is

A linear covariate can absorb only a first-order quality effect, while gene detection depends on completeness non-linearly, and the negative control shows the expected trace of that: across the 92 ribosomal orthologues, carriage rate correlates with the within-genus effect at *r* = +0.314, the direction under-correction produces. We therefore tested non-linear covariates directly, adding squared, cubed and piecewise-linear terms in completeness and feature count to the residualisation, and re-ran the main contrast on nested completeness subsets (≥ 70%, ≥ 90%, ≥ 95%, ≥ 97%).

The non-linear terms tighten the negative control substantially without touching the signal. Adding squared terms moves the artefact floor, the 95th percentile of |controlled effect| across the ribosomal set, from 0.055 to 0.031 and the ribosomal maximum from 0.072 to 0.043, while every large surviving effect holds or grows: alsT +0.231 to +0.241, betT +0.163 to +0.189, SLC13 +0.151 to +0.168, phr +0.156 to +0.158, phrB +0.191 to +0.180. Cubic terms give 0.031 with the same signals; a completeness spline gives 0.052. A specification that removed real signal along with the artefact would collapse both columns together, and this one does not, which identifies the residue as assembly non-linearity rather than biology (Extended Data Fig. 7). We nevertheless report the linear specification throughout, because it is the conservative choice: it leaves the artefact floor roughly twice as high, so every feature we report clears a threshold that a better-specified model would have lowered. Readers who prefer the quadratic specification should read our survivor set as an underestimate. Retention is 0.097, 0.117, 0.112 and 0.142 and the inflation factor 8.4, 7.9, 8.4 and 6.7, both stable, with no trend that would indicate the linear covariates are failing, while the mean ribosomal *t* falls from 0.554 to 0.26, 0.24 and 0.29. The survivor count falls from 59 to 28, 18 and 13 across the same subsets, but that is sample size: the number of paired genera falls from 75 to 26 at the same time, so the count is not evidence either way.

The residue that remains on the negative control under every specification, a mean *t* of +0.55 against a true value of zero, is assembly incompleteness rather than biology. Holding the number of genomes, the genus composition and the two-sided balance of each genus exactly fixed, and varying only *which* genomes are drawn, the most complete in each cell, a random draw, or the least complete, moves it monotonically with assembly quality: mean ribosomal *t* is +0.67 at 91.9% mean completeness, +0.46 to +0.53 at 93.5%, +0.24 at 95.1% and +0.18 at 95.9% (r = −0.986, slope −0.133 per completeness point; Extended Data Fig. 1c). An earlier attempt to show this failed because tightening quality also changed sample size and the two effects cancelled. The same phenomenon is the RNA-polymerase and ribosomal tail in the isolate-versus-MAG comparison, where it appears at its largest because isolate and MAG assemblies differ most (Extended Data Fig. 6b).

This also disposes of the one whole-proteome trait that would otherwise embarrass the analysis. Mean protein length retains 0.87, higher than any trait anyone has proposed as adaptive, which would undercut any reading of retention as evidence of adaptation. It is not biology: mean protein length correlates with completeness at r = +0.53 (*P* = 10 ¹²), with a slope of 1.77 residues per completeness point, and ocean genomes are 3.98 completeness points more complete than soil genomes on average, which alone predicts a 7.0-residue difference, larger than the 2.1 residues actually observed. Fragmented assemblies truncate open reading frames, and this trait reads that out rather than the proteome. It is therefore excluded from the whole-proteome comparison.

### Phylogenetic mixed model

25 ribosomal proteins located by KEGG gene-level annotation, aligned with MAFFT v7.520^36^, columns with > 50% gaps removed (3,925 columns), tree inferred with FastTree 2.2.0^37^ under LG+Γ. Kinship is the Gower-centred^43^ patristic distance matrix, centred on the analysis set; the variance-component formulation is the one standard in association studies with sample structure^45^. The variance ratio is estimated by REML on a grid and shared across features; because it is not identified, results are reported across the grid. Per-feature h² is the per-feature REML optimum on a fine grid. Comparisons between this model and the main analysis are made under matched specifications, per-genus covariate slopes in both, so that the ladder measures phylogeny rather than the specification change.

### Within-species analysis

To test the scale below the genus we matched genomes by GTDB species name across the soil and ocean catalogues. Only labels resolved to a binomial were used; SMAG records an unassigned species as the genus name alone, and treating those as species would create false matches at genus level. Because both catalogues are dereplicated to species representatives, no species has more than two genomes on either side and no species has three on both, so the within-genus design cannot be repeated at this level. What is possible is a paired design: 73 species contribute one soil and one ocean representative each (74 pairs). Per-feature effects are the mean of the 74 paired differences. A stratum of two genomes cannot support a within-stratum covariate fit, so quality was instead adjusted across pairs, regressing the 74 difference vectors on the per-pair differences in completeness and contamination and retaining the intercept. We report the ribosomal control at this level as the criterion for whether any of it is interpretable.

### Matched-null test for published claims

For each claim, random gene sets of the same size were drawn from the catalogue and matched on |naive Δ|; stricter nulls additionally drew only from KOs assigned to a KEGG pathway and matched each member on carriage rate. Because within-genus residualisation is linear, a composite feature’s effect equals the mean of its members’ effects, so null sets are assembled from per-KO coefficients. The test is performed at panel level: one matched set per claim, panel median recorded, 5,000 repetitions, *P* = the fraction of null panels reaching the observed median. The null value quoted against the observed one is therefore the median over those 5,000 panel medians, 0.089, and not the median of the 23 per-claim nulls, 0.091, which Extended Data Table 1 lists and which a reader recomputing from that table would obtain; the two aggregations differ by 0.002 and only the first is the quantity the *P* value is computed from. Naive thresholds of 0.02, 0.10 and 0.20 define nested subsets and are three levels of one test, not three independent confirmations. Because the panel was assembled by us rather than sampled by a pre-registered protocol, we additionally report a leave-one-claim-out analysis: each claim is dropped in turn and the panel test re-run, and we report the worst-case p rather than only the full-panel p.

Claim-panel effects are reported under the three-covariate main specification, so that the panel is judged by the same standard as everything else in this paper. A second estimate exists because several panel members are drawn from Pfam and the two feature spaces have to be estimated the same way for a panel median to mean anything, which argues for residualising on feature count alone. The choice does not carry the result: on feature count alone the 23 claims retain 0.246 against a matched-null 0.103 (2.4-fold, *P* = 0.0008; leave-one-claim-out worst case *P* = 0.0016), against 0.237 versus 0.089 (2.7-fold, *P* = 0.0018, worst case 0.0028) under the specification reported. Every claim is listed under both in Extended Data Table 1.

### Held-out-genus cross-validation and feature ablation

L2-regularised logistic regression (scikit-learn 1.6.1^38^, C = 0.05) predicting biome from binary feature vectors, with entire genera held out across eight folds so that no genus appears in both training and test. The ablation asks whether the features a per-feature test selects are where the classifier’s information is. Deleting a set of features and observing a drop is uninterpretable on its own, because deleting any features at all changes the problem; every deletion is therefore paired with a matched random deletion, the same number of features, drawn to match the deleted set’s carriage distribution in twenty prevalence bins, repeated 20 times, with the mean and standard deviation of the resulting AUC reported. We also report a stepwise curve in which the top *k* features by |t| are deleted for *k* = 64, 128, …, 4,096, each against its own matched random deletion; Fig. 4b plots the curve to *k* = 4,096. Beyond *k* = 1,024 more than an eighth of the matrix is gone and the deletion is no longer a statement about a feature set, so the curve is read only to that point. An earlier version of this analysis deleted only the six KEGG features (five Pfam) passing a since-abandoned magnitude criterion and reported the resulting negligible AUC change as evidence that screening misses the signal; that conclusion does not survive the corrected ablation and has been withdrawn.

### Nested cross-validation of the ablation

A matched random deletion controls for the size and carriage of the deleted set but not for how the set was chosen. The significant features are selected by a test fitted to every genus, including the genera the classifier is then tested on, whereas the random features have seen no labels at all; part of any difference between them is therefore selection reaching the test fold rather than information carried by the features. We remove this by refitting the selection inside each fold. For each of the eight held-out-genus folds, the within-genus contrast and the Westfall–Young threshold are re-estimated on that fold’s training genera alone, the features that fold selects are deleted from both training and test matrices, and the classifier is trained and its held-out predictions recorded. Deletion sets therefore differ between folds, but every prediction is out of sample with respect to both the classifier and the feature selection, so the pooled AUC is a valid estimate of what a genuinely prospective screen would cost. Both the selection-on-everything and the nested figure are reported, the first as an upper bound.

### The contrast series and the taxonomic-overlap relationship

Eight biome pairs were run through one pipeline on Pfam features (present in ≥ 50 genomes of the pair): soil–ocean, soil–lake, soil–river, lake– ocean, river–ocean, lake–river, gut–soil and gut–ocean. For each we record the genus-level Jaccard overlap between the two catalogues, the fraction of features reaching naive |Δ| ≥ 0.20, and the median retention among those features, and relate the first to the other two by Spearman correlation. Lake–river carries no environmental contrast and serves as a negative control at the level of the design: both catalogues are freshwater and were assembled independently by different groups, so whatever a conventional comparison reports between them is catalogue batch effect and lineage composition alone. The eight contrasts are not mutually independent, two share the gut catalogue and four share soil or ocean, so the correlation is reported with that caveat and without an interval.

### Choice of estimator

The within-lineage contrast is the estimator bacterial association studies standardised, used deliberately so that the inflation we report cannot be attributed to a new method. On the phylogeny subset a single-kinship mixed model of the kind those studies fit passes 36 features, where a two-axis principal-coordinates correction passes 21: the standard correction is the more permissive of the two, so the collapse is not the product of a control chosen for being severe.

### Comparison with the within-species regime

To place our contrast against the setting in which correcting for population structure is already standard, we re-analysed the association data of ref. 21, six host-niche contrasts in a collection of *Escherichia coli* isolated from digestive tracts, using our own estimator rather than reading their coefficients: what they report is a structure-corrected logistic coefficient, which is not the quantity our retention ratio divides. For each of their runs we rebuilt both arms on the carriage-rate scale from the carrier and non-carrier lists their output records together with their phenotype table, took lineage strata to be average-linkage clusters of their own distance matrix, and estimated the within-stratum contrast by the same residualisation we apply within genera. Lineage imbalance is the share of phenotype variance those strata explain. Retention is the median of per-variant ratios over the variants their filter passed, so it is the analogue of our figure for features a conventional analysis calls strongly enriched rather than of a genome-wide one, and our own contrast is recomputed on that same statistic for the comparison, giving 0.288 against their 0.918 to 1.003.

Their 100 runs per contrast are subsamples of one strain collection, and the contrasts share most of their genomes with each other, so the unit of analysis is the six contrasts and we do not report a correlation across runs. Two controls bound the comparison. Retention and imbalance are computed from the same strata, so part of any association between them could be mechanical; partialling out how stratum-structured each variant is leaves it unchanged. Stratification granularity also differs between the two settings, so the analysis is repeated at four target strata counts spanning 39 to 16 genomes per stratum, which brackets our 22: retention in their data is 0.99, 0.96, 0.91 and 0.91 across that range, so the gap is not an artefact of how finely lineage is cut.

### Whole-proteome traits

Ten traits computed over all proteins of each genome: isoelectric point, the fraction of acidic proteins, acidic, basic and total charged residue fractions, elemental economy (N and C atoms per residue), aromatic residue fraction, GRAVY hydrophobicity and mean protein length. Mean protein length is excluded for the reason given below, so nine enter the comparison. These traits share no methodology with the gene-content analysis, which is the point of including them, but they are not immune to assembly quality: mean protein length in particular is sensitive to contig fragmentation, which truncates ORFs, and is excluded for the reason given above. Across all proteins of the paired genomes, overall isoelectric point differs by −0.140 naively (soil 6.502, ocean 6.362) but by only −0.028 within genera (retention 0.20, *P* = 0.12); nitrogen atoms per residue retain 0.27 (*P* = 0.02); GRAVY hydrophobicity retains 0.06 (*P* = 0.08). The published claim that marine proteomes are systematically more acidic^46^ is directionally correct but retains only 20% of its magnitude within genera and is not significant there (Extended Data Fig. 3).

### Claim panel

27 adaptation claims drawn from the literature^46,48–73^, each mapped to KOs verified through the KEGG REST API; 23 were auditable in our data at naive |Δ| ≥ 0.02. Extended Data Table 1 lists all 27 with their source, feature set, naive and controlled effects, retention, matched-null percentile and auditability, together with the type of evidence on which the original claim rested (genome prevalence, metagenomic read abundance, transcript abundance, biochemical or physiological characterisation, or phylogenetic reconstruction). Because a claim originally made on read abundance and a claim originally made on genome prevalence are not the same quantity, retention is reported for the prevalence-based subset alone as well as for the whole panel: 6 of the 23 auditable claims rest on genome prevalence and retain a median 0.18, against 0.24 for all 23 and 0.34 for the 17 resting on other evidence, so the re-test is if anything harsher on the claims measured in our own quantity. The classification records the design the source paper reports; for four of the panel’s 26 source references we confirmed it against the paper directly, and for the remainder it is taken from the stated design, which the table marks. Retention ratio is the controlled effect divided by the naive effect. The panel is a representative selection of recurring claims, not a systematic-review sample, and we do not present it as one; the leave-one-claim-out analysis above is the check on whether any single choice carries the result.

## Figure legends

Unless stated otherwise, all statistical tests are two-sided; the analysis unit is the genome, the resampling and exchangeability unit is the genus, and effect sizes are differences in carriage rate (the proportion of genomes carrying a feature). “Controlled” means the within-genus estimate under the main specification, residualisation on annotated feature count, CheckM completeness and CheckM contamination with per-genus slopes. The soil–ocean contrast comprises 1,682 genomes in 75 paired genera and 8,077 KEGG orthologues (7,706 Pfam domains, 78 paired genera). Confidence intervals are percentile intervals from 200 bootstrap resamples of genera; the naive contrast is a whole-catalogue quantity and is held fixed across resamples, so intervals are shown only for quantities that depend on the within-genus layer.

## Author contributions

K.Z. created the study design. K.Z., Y.W. and R.X. collected all datasets. K.Z. and R.X. performed the data analysis and visualization. K.Z., Y.W. and R.X. contributed to the scientific discussion and wrote the manuscript. All authors read and approved the final manuscript.

## References

1. Baas Becking, L. G. M. Geobiologie of Inleiding tot de Milieukunde (W. P. van Stockum & Zoon, 1934).

2. de Wit, R. & Bouvier, T. ‘Everything is everywhere, but, the environment selects’; what did Baas Becking and Beijerinck really say? Environ. Microbiol. 8, 755–758 (2006).

3. O’Malley, M. A. The nineteenth century roots of ‘everything is everywhere’. Nat. Rev. Microbiol. 5, 647–651 (2007).

4. Martiny, J. B. H. et al. Microbial biogeography: putting microorganisms on the map. Nat. Rev. Microbiol. 4, 102–112 (2006).

5. Price, A. L. et al. Principal components analysis corrects for stratification in genome-wide association studies. Nat. Genet. 38, 904–909 (2006).

6. Brynildsrud, O., Bohlin, J., Scheffer, L. & Eldholm, V. Rapid scoring of genes in microbial pan-genome-wide association studies with Scoary. Genome Biol. 17, 238 (2016).

7. Collins, C. & Didelot, X. A phylogenetic method to perform genome-wide association studies in microbes that accounts for population structure and recombination. PLoS Comput. Biol. 14, e1005958 (2018).

8. Earle, S. G. et al. Identifying lineage effects when controlling for population structure improves power in bacterial association studies. Nat. Microbiol. 1, 16041 (2016).

9. Lees, J. A., Galardini, M., Bentley, S. D., Weiser, J. N. & Corander, J. pyseer: a comprehensive tool for microbial pangenome-wide association studies. Bioinformatics 34, 4310–4312 (2018).

10. Jaillard, M. et al. A fast and agnostic method for bacterial genome-wide association studies: bridging the gap between k-mers and genetic events. PLoS Genet. 14, e1007758 (2018).

11. Göring, H. H. H., Terwilliger, J. D. & Blangero, J. Large upward bias in estimation of locus-specific effects from genomewide scans. Am. J. Hum. Genet. 69, 1357–1369 (2001).

12. Ioannidis, J. P. A. Why most discovered true associations are inflated. Epidemiology 19, 640–648 (2008).

13. Louca, S., Parfrey, L. W. & Doebeli, M. Decoupling function and taxonomy in the global ocean microbiome. Science 353, 1272–1277 (2016).

14. Louca, S. et al. High taxonomic variability despite stable functional structure across microbial communities. *Nat*. Ecol. Evol. 1, 0015 (2016).

15. Martiny, J. B. H., Jones, S. E., Lennon, J. T. & Martiny, A. C. Microbiomes in light of traits: a phylogenetic perspective. Science 350, aac9323 (2015).

16. Martiny, A. C., Treseder, K. & Pusch, G. Phylogenetic conservatism of functional traits in microorganisms. ISME J. 7, 830–838 (2013).

17. Maistrenko, O. M. et al. Disentangling the impact of environmental and phylogenetic constraints on prokaryotic within-species diversity. ISME J. 14, 1247–1259 (2020).

18. Douglas, G. M., Hayes, M. G., Langille, M. G. I. & Borenstein, E. Integrating phylogenetic and functional data in microbiome studies. Bioinformatics 38, 5055–5063 (2022).

19. Manor, O. & Borenstein, E. Systematic characterization and analysis of the taxonomic drivers of functional shifts in the human microbiome. Cell Host Microbe 21, 254–267 (2017).

20. Tamames, J., Sánchez, P. D., Nikel, P. I. & Pedrós-Alió, C. Quantifying the relative importance of phylogeny and environmental preferences as drivers of gene content in prokaryotic microorganisms. Front. Microbiol. 7, 433 (2016).

21. Monteith, W. et al. Everything is everywhere but *Escherichia coli* adapts to different niches. ISME J. 20, wraf267 (2026).

22. Ma, B. et al. A genomic catalogue of soil microbiomes boosts mining of biodiversity and genetic resources. Nat. Commun. 14, 7318 (2023).

23. Chen, J. et al. Global marine microbial diversity and its potential in bioprospecting. Nature 633, 371–379 (2024).

24. Garner, R. E. et al. A genome catalogue of lake bacterial diversity and its drivers at continental scale. Nat. Microbiol. 8, 1920–1934 (2023).

25. Borton, M. A. et al. A functional microbiome catalogue crowdsourced from North American rivers. Nature 637, 103–112 (2025).

26. Almeida, A. et al. A unified catalog of 204,938 reference genomes from the human gut microbiome. Nat. Biotechnol. 39, 105–114 (2021).

27. Dai, R. et al. Crop root bacterial and viral genomes reveal unexplored species and microbiome patterns. Cell 188, 2521–2539.e22 (2025).

28. Cantalapiedra, C. P., Hernández-Plaza, A., Letunic, I., Bork, P. & Huerta-Cepas, J. eggNOG-mapper v2: functional annotation, orthology assignments, and domain prediction at the metagenomic scale. Mol. Biol. Evol. 38, 5825–5829 (2021).

29. Huerta-Cepas, J. et al. eggNOG 5.0: a hierarchical, functionally and phylogenetically annotated orthology resource. Nucleic Acids Res. 47, D309–D314 (2019).

30. Kanehisa, M. & Goto, S. KEGG: Kyoto Encyclopedia of Genes and Genomes. Nucleic Acids Res. 28, 27–30 (2000).

31. Mistry, J. et al. Pfam: the protein families database in 2021. Nucleic Acids Res. 49, D412–D419 (2021).

32. Parks, D. H., Imelfort, M., Skennerton, C. T., Hugenholtz, P. & Tyson, G. W. CheckM: assessing the quality of microbial genomes recovered from isolates, single cells, and metagenomes. Genome Res. 25, 1043–1055 (2015).

33. Parks, D. H. et al. GTDB: an ongoing census of bacterial and archaeal diversity through a phylogenetically consistent, rank normalized and complete genome-based taxonomy. Nucleic Acids Res. 50, D785–D794 (2022).

34. Chaumeil, P.-A., Mussig, A. J., Hugenholtz, P. & Parks, D. H. GTDB-Tk v2: memory friendly classification with the genome taxonomy database. Bioinformatics 38, 5315–5316 (2022).

35. Bowers, R. M. et al. Minimum information about a single amplified genome (MISAG) and a metagenome-assembled genome (MIMAG) of bacteria and archaea. Nat. Biotechnol. 35, 725–731 (2017).

36. Katoh, K. & Standley, D. M. MAFFT multiple sequence alignment software version 7: improvements in performance and usability. Mol. Biol. Evol. 30, 772–780 (2013).

37. Price, M. N., Dehal, P. S. & Arkin, A. P. FastTree 2, approximately maximum-likelihood trees for large alignments. PLoS ONE 5, e9490 (2010).

38. Pedregosa, F. et al. Scikit-learn: machine learning in Python. J. Mach. Learn. Res. 12, 2825–2830 (2011).

39. Westfall, P. H. & Young, S. S. Resampling-Based Multiple Testing: Examples and Methods for p-Value Adjustment (Wiley, 1993).

40. Benjamini, Y. & Hochberg, Y. Controlling the false discovery rate: a practical and powerful approach to multiple testing. J. R. Stat. Soc. B 57, 289–300 (1995).

41. Frisch, R. & Waugh, F. V. Partial time regressions as compared with individual trends. Econometrica 1, 387–401 (1933).

42. Lovell, M. C. Seasonal adjustment of economic time series and multiple regression analysis. J. Am. Stat. Assoc. 58, 993–1010 (1963).

43. Gower, J. C. Some distance properties of latent root and vector methods used in multivariate analysis. Biometrika 53, 325–338 (1966).

44. Cameron, A. C., Gelbach, J. B. & Miller, D. L. Bootstrap-based improvements for inference with clustered errors. Rev. Econ. Stat. 90, 414–427 (2008).

45. Kang, H. M. et al. Variance component model to account for sample structure in genome-wide association studies. Nat. Genet. 42, 348–354 (2010).

46. Jurdzinski, K. T. et al. Large-scale phylogenomics of aquatic bacteria reveal molecular mechanisms for adaptation to salinity. Sci. Adv. 9, eadg2059 (2023).

47. Tito, R. Y. et al. Microbiome confounders and quantitative profiling challenge predicted microbial targets in colorectal cancer development. Nat. Med. 30, 1339–1348 (2024).

48. Béjà, O. et al. Bacterial rhodopsin: evidence for a new type of phototrophy in the sea. Science 289, 1902–1906 (2000).

49. Galinski, E. A. & Trüper, H. G. Microbial behaviour in salt-stressed ecosystems. FEMS Microbiol. Rev. 15, 95–108 (1994).

50. Sinha, R. P. & Häder, D.-P. UV-induced DNA damage and repair: a review. Photochem. Photobiol. Sci. 1, 225–236 (2002).

51. Villarreal-Chiu, J. F., Quinn, J. P. & McGrath, J. W. The genes and enzymes of phosphonate metabolism by bacteria, and their distribution in the marine environment. Front. Microbiol. 3, 19 (2012).

52. Luo, H., Benner, R., Long, R. A. & Hu, J. Subcellular localization of marine bacterial alkaline phosphatases. Proc. Natl Acad. Sci. USA 106, 21219–21223 (2009).

53. Howard, E. C. et al. Bacterial taxa that limit sulfur flux from the ocean. Science 314, 649–652 (2006).

54. Moran, M. A. et al. Genomic insights into bacterial DMSP transformations. Ann. Rev. Mar. Sci. 4, 523–542 (2012).

55. Noinaj, N., Guillier, M., Barnard, T. J. & Buchanan, S. K. TonB-dependent transporters: regulation, structure, and function. Annu. Rev. Microbiol. 64, 43–60 (2010).

56. Cordero, P. R. F. et al. Atmospheric carbon monoxide oxidation is a widespread mechanism supporting microbial survival. ISME J. 13, 2868–2881 (2019).

57. Bayer, B. et al. Ammonia-oxidizing archaea release a suite of organic compounds potentially fueling prokaryotic heterotrophy in the ocean. Environ. Microbiol. 21, 4062–4075 (2019).

58. Berlemont, R. & Martiny, A. C. Phylogenetic distribution of potential cellulases in bacteria. Appl. Environ. Microbiol. 79, 1545–1554 (2013).

59. Bai, Y. et al. Functional overlap of the *Arabidopsis* leaf and root microbiota. Nature 528, 364– 369 (2015).

60. Beier, S. & Bertilsson, S. Bacterial chitin degradation, mechanisms and ecophysiological strategies. Front. Microbiol. 4, 149 (2013).

61. Galperin, M. Y., Mekhedov, S. L., Puigbo, P., Smirnov, S., Wolf, Y. I. & Rigden, D. J. Genomic determinants of sporulation in Bacilli and Clostridia. Environ. Microbiol. 14, 2870–2890 (2012).

62. Pitcher, R. S., Brissett, N. C. & Doherty, A. J. Nonhomologous end-joining in bacteria: a microbial perspective. Annu. Rev. Microbiol. 61, 259–282 (2007).

63. Wattam, A. R. et al. Improvements to PATRIC, the all-bacterial bioinformatics database and analysis resource center. Nucleic Acids Res. 45, D535–D542 (2017).

64. Raymond, J., Siefert, J. L., Staples, C. R. & Blankenship, R. E. The natural history of nitrogen fixation. Mol. Biol. Evol. 21, 541–554 (2004).

65. Nies, D. H. Efflux-mediated heavy metal resistance in prokaryotes. FEMS Microbiol. Rev. 27, 313–339 (2003).

66. Vasu, K. & Nagaraja, V. Diverse functions of restriction-modification systems in addition to cellular defense. Microbiol. Mol. Biol. Rev. 77, 53–72 (2013).

67. Makarova, K. S. et al. Evolutionary classification of CRISPR–Cas systems: a burst of class 2 and derived variants. Nat. Rev. Microbiol. 18, 67–83 (2020).

68. Tabita, F. R., Hanson, T. E., Satagopan, S., Witte, B. H. & Kreel, N. E. Phylogenetic and evolutionary relationships of RubisCO and the RubisCO-like proteins. Philos. Trans. R. Soc. B 363, 2629–2640 (2008).

69. Häse, C. C. & Barquera, B. Role of sodium bioenergetics in *Vibrio cholerae*. Biochim. Biophys. Acta 1505, 169–178 (2001).

70. Wadhams, G. H. & Armitage, J. P. Making sense of it all: bacterial chemotaxis. Nat. Rev. Mol. Cell Biol. 5, 1024–1037 (2004).

71. Fuchs, G., Boll, M. & Heider, J. Microbial degradation of aromatic compounds, from one strategy to four. Nat. Rev. Microbiol. 9, 803–816 (2011).

72. Martiny, A. C., Coleman, M. L. & Chisholm, S. W. Phosphate acquisition genes in *Prochlorococcus* ecotypes: evidence for genome-wide adaptation. Proc. Natl Acad. Sci. USA 103, 12552–12557 (2006).

73. Ghosh, W. & Dam, B. Biochemistry and molecular biology of lithotrophic sulfur oxidation by taxonomically and ecologically diverse bacteria and archaea. FEMS Microbiol. Rev. 33, 999– 1043 (2009).

