## SUPPLEMENTARY for "Lineage sorting inflates cross-environment functional enrichment"

**Supplementary Table 1 | A checklist for cross-environment functional enrichment**

Every item costs analysis time rather than new sequencing, and every item is one we got wrong in some draft of this work before the diagnostic named in the third column caught it. Items 1 to 3 are the three the Discussion names as required; items 4 to 13 are the rest of what our own audit turned up. The point of the “evidence” column is that none of these are stylistic preferences: each one is a place where the uncorrected and corrected answers differed by enough to change a conclusion.

| **#** | **Do this** | **Instead of** | **Why: evidence from this study** | **What it costs** |
| --- | --- | --- | --- | --- |
| **1** | **Estimate the contrast within a lineage** held fixed at a comparable rank (GTDB genus, whose boundaries are set by relative evolutionary divergence), using only lineages present on both sides | Comparing whole catalogues across environments | Median effect of the 178 orthologues called strongly enriched fell from 0.250 to 0.030; regressing controlled on naive effect across all 8,077 gave a slope of 0.109, so a reported 30-percentage-point difference is about three within a genus. 45 of 178 reversed sign | Restricts the analysis to shared lineages: 1,682 of 22,905 genomes, 75 of several thousand genera. This is the real cost and it is large |
| **2** | **Compute an artefact floor per contrast and per carriage band**: the 95th percentile of \|controlled effect\| across genes whose true effect is zero, with a bootstrap interval, and separately by permuting the environment label within lineage and taking the 95th percentile within bins of carriage. Require a reported feature to clear the larger of the two in its own band | One global magnitude threshold, or none | The floor ran from 0.031 (CRBC to ocean) to 0.094 (CRBC to soil) across biome contrasts and 0.219 at the species level, a sevenfold range; a magnitude interpretable in one contrast is noise in another. The band matters as much as the contrast: single-copy genes cannot be found between 10% and 40% carriage, which is where most of our survivors sit, and where they can be found, at 40 to 80%, their own percentile is 0.059 against a whole-set scalar of 0.055 | One extra estimation pass over ~100 null genes, plus one permutation pass |
| **3** | **Set the significance threshold by permutation** of the environment label within lineage, taking the maximum standardised statistic across all features (Westfall–Young), and report its Monte-Carlo error | A magnitude cut-off such as \|Δ\| ≥ 0.15, or per-feature FDR alone | A magnitude cut-off discards 48 of our 59 survivors, and half the survivors have naive effects too small to enter a conventional list at all. The threshold is also under-resampled easily: at 200 permutations the survivor count moves by roughly ±10, so we use 10,000 | 10,000 permutations of the full design |
| **4** | **Control assembly quality inside each stratum**, with per-lineage covariate slopes, on completeness, contamination and annotated feature count | Pooling covariate slopes, or filtering on quality instead of modelling it | Pooled slopes carry twice the residual bias on the negative control (mean *t* +1.03 against +0.55). Raising the quality filter instead removes genomes non-randomly with respect to biome and lineage | Degrees of freedom in small strata; strata too small to fit are centred only |
| **5** | **Carry an internal negative control** of genes whose true between-environment difference is zero, and report its median, its upper tail and its mean *t* | Reporting only that the method “controls for” quality | Without quality covariates the largest within-genus effects in our dataset were ribosomal proteins. With them the ribosomal median went 0.020 to 0.007 but the mean *t* stayed at +0.55, a detectable residual bias we could then diagnose and explain rather than conceal | Nothing; the genes are already in the matrix |
| **6** | **Test whether the residue is non-linear** by adding squared or spline terms in completeness, and check the null genes and the surviving signal separately | Assuming a linear covariate has absorbed the quality effect | Squared terms moved the single-copy floor from 0.055 to 0.031 while every large surviving effect held or grew. Both columns falling together would have meant over-correction; only the null falling means correction. We kept the linear specification anyway, as the one that leaves the higher floor (Extended Data Fig. 7) | One extra specification |
| **7** | **Measure taxonomic overlap before testing any gene, and treat it as an ordering rather than a prediction.** Low genus-level overlap between the two catalogues goes with a larger fraction of the genome being called enriched | Assuming the size of the problem is a constant, or treating the association as a calibration | The fraction of features called strongly enriched ran from 0.08% to 20% across eight contrasts, more than two orders of magnitude, and tracked overlap (Spearman rho = −0.71, *P* = 0.047). But the eight contrasts share catalogues, so *n* = 8 overstates the degrees of freedom, and by the standard we applied to a similar relationship in Note 1 this licenses an expectation about direction, not a predicted value. Our main contrast, at 3%, is a point on that range rather than a typical value | One taxonomy intersection |
| **8** | **Resample the exchangeable unit** (lineages) for intervals, relabelling so a lineage drawn twice contributes two independent strata | Bootstrapping genomes | Genomes within a lineage are not exchangeable with each other, so a genome-level bootstrap understates the interval. Check the bootstrap estimator reproduces the point fit on unresampled data before trusting it | 200 resamples |
| **9** | **Re-annotate every catalogue through one pipeline from protein sequence**, and sanity-check a universal gene before any comparison | Comparing catalogues’ published annotations | Under UHGG’s native annotations RecA, universal by definition, appears in 0.0% of gut genomes, because Pfam had renamed the domain RecA_N. Accession matching recovers 96.3% and re-annotation 95.8%, while ribosomal domains match under either scheme, so the failure is both silent and selective | One annotation run per catalogue |
| **10** | **Report at the level you tested.** If features were tested individually, do not aggregate to pathways or modules and present that as a separate result | Reporting module- or pathway-level enrichment as independent corroboration | With a linear estimator a module effect is exactly the mean of its member genes’ effects, so it is a deterministic consequence, not a second measurement. 34 of 239 modules reached \|Δ\| ≥ 0.05 conventionally and none survived, because averaging one or two real signals against dozens of hitchhikers is arithmetic | Nothing; it removes an analysis |
| **11** | **Pair every feature-deletion or ablation argument with a matched random deletion** of the same size and carriage distribution, and re-select the deleted set inside each cross-validation fold | Deleting a selected feature set and reporting the AUC drop | Deleting our significant set cost 0.234 of AUC with selection fitted on all genera and 0.080 with selection refitted inside each fold, so two-thirds of the apparent effect was selection reaching the test fold. An earlier version of this analysis reported a conclusion in the opposite direction and it did not survive | One nested cross-validation |
| **12** | **Include a design-level negative control**: two independently built catalogues from the *same* environment, so any apparent enrichment between them is batch effect and lineage composition alone | Relying only on gene-level negative controls | Lake against river, both freshwater and built by different groups, had the highest genus overlap of any pair we ran and a conventional analysis called 0.08% of features strongly enriched. Had our collapse been an artefact of comparing two independently built catalogues, it would have appeared here | One extra contrast, if a suitable pair exists |
| **13** | **State whether “enrichment” means carriage or abundance**, and check the exchangeability of the permutation unit by restricting permutation to sub-clades | Leaving the quantity implicit; permuting freely within lineage | Between-environment patristic distances exceed within-environment distances in 39 of 58 genera, making free within-lineage permutation mildly anti-conservative; restricting to sub-clades at a stricter cut-off reduced the survivor count by 29%, which is why we report a range rather than a point. Carriage is also the conservative quantity: an abundance-weighted comparison lets one dominant lineage contribute its whole gene complement | A tree for the analysis set |
| **14** | **Check where the point estimate falls among your bootstrap replicates** before quoting a percentile interval, and switch to estimate ±1.96 bootstrap s.e. or a bias-corrected interval when it does not sit near the middle | Quoting the 2.5th and 97.5th percentiles of the replicates by default | Our headline statistic is a median of absolute values. Resampling lineages injects noise, \|·\| is inflated by noise, and the replicate distribution therefore sits above the point estimate, which lands at its 12.5th percentile; the percentile interval was 5:1 lopsided about the estimate it was supposed to bracket, while the slope and retention statistics from the same bootstrap were centred and unaffected | One line: the quantile of the estimate within the replicates |

**How to read a published enrichment result you cannot re-analyse**

Five of the items above can be applied to someone else’s paper from its methods section alone.

1. **Does the paper report an uncorrected and a corrected estimate for the same features?** If it reports only one, no inflation figure can be recovered from it, whatever the correction. A partitioned variance ratio, a consistency test across lineages and a niche-versus-niche comparison all lack the uncorrected arm.
2. **Was the comparison within lineage?** If not, treat the reported effect sizes as upper bounds. For an individual gene our best estimate of what remains is about a tenth of the reported value; for a claim assembled from several well-characterised genes, about a quarter.
3. **What was the taxonomic overlap between the two sets being compared?** The lower it is, the larger the fraction of the genome a conventional analysis will call enriched. Comparisons across very different environments, where overlap is lowest, are the ones whose enrichment lists are least trustworthy, which is the opposite of the intuition that a big biological difference makes a result safer.
4. **Was the threshold a magnitude cut-off?** If so, the reported set is biased towards widely carried genes and against exactly the small, real, within-lineage effects that survive lineage control.
5. **Which regime is the study in?** Ancestry correction is standard in bacterial association studies, and in that setting it barely changes the answer: applying our estimator to six host-niche contrasts within one *Escherichia coli* collection retains 92 to 100% of the uncorrected effect and reverses no signs. A study comparing strains of one species across niches is in that regime and its uncorrected effect sizes are roughly right. A study comparing catalogues built from different environments is in ours, where the same estimator leaves 29% and reverses a quarter of signs. The question is not whether the authors corrected but which of the two situations they were in.

None of this implies the underlying findings are false. Our audited claims beat matched random gene sets by 2.7-fold in retention and 2.4-fold in controlled effect: the field has been identifying real signal and mis-attributing its cause.

**Supplementary Note 1 | Analyses withdrawn during this work**

We list these because each was, at some point, a result we believed, and because the diagnostic that killed each one is reusable.

| **Withdrawn claim** | **Why it failed** | **Diagnostic that caught it** |
| --- | --- | --- |
| A photolyase enrichment on the rhizosphere contrast | Fell below that contrast’s own artefact floor | Per-contrast artefact floor (item 2) |
| “Deleting every significant feature barely changes classifier AUC, so screening misses the signal” | The deletion was not paired with a matched random deletion; with one, and with selection refitted out of sample, the cost is 0.080, not negligible | Matched random deletion plus nested selection (item 11) |
| A species-level directional result, 39 of 59 survivors keeping sign | 13 of the 59 had within-species effects of exactly zero, whose sign was floating-point residue; removing them left the effect entirely in the ocean-positive half, which a directional completeness gap of +3.7 percentage points would inflate, and it vanished in completeness-matched pairs | Sign retention split by side, plus a quality-matched subset (Extended Data Fig. 8) |
| Mean protein length as an adaptive whole-proteome trait, retention 0.87 | Correlates with completeness at *r* = +0.53; the completeness gap between catalogues alone predicts a larger difference than is observed | Trait against assembly quality (Extended Data Fig. 3c) |
| “The inflation factor is the same at every analytical level” | The whole-proteome value supporting it could not be reproduced from the analysis outputs; the correct median is 0.02, not 0.16, so retention varies about tenfold across levels | Deriving figure values from the output tables rather than typing them in |
| A pathway-level retention of 0.09, quoted in the text and drawn in Fig. 5a | No estimator on the pathway output table reproduces it; the median signed retention over the 239 modules, computed as at every other level, is 0.06 | Deriving every figure value from the output tables rather than typing them in, and a scalar check that fails when text and table disagree |
| “Retention is a predictable function of lineage imbalance, so the size of the problem can be calibrated in advance” | The relationship is real within one collection but its 564 runs are 100 subsamples of each of six contrasts that share most of their genomes; at the honest unit the correlation is rho = -0.66 on n = 6. Our own contrast also sits far off the line, retaining 29% where it predicts 83% | Checking sample overlap between runs before trusting a correlation across them |
| “The artefact floor calibrated on single-copy genes is roughly twice as strict as a prevalence-matched one” | True against a permutation null, which contains no bias term, but not against single-copy genes restricted to the survivors’ own carriage band, where the percentile is 0.059 against the scalar’s 0.055 | Comparing like with like: a null that carries the bias against a null that carries the bias |

**Supplementary Note 2 | What the design cannot do**

- **Adaptation fixed deep in the tree is invisible to it.** Anything that sorted before the genera we compare existed is absorbed into the lineage term and counted as sorting. This is a limit of the question, not of the estimator, and it means our within-lineage layer is a lower bound on environmental adaptation in total.
- **Genera shared across environments are cosmopolitan and probably generalist.** That bias runs towards finding within-lineage adaptation, so it strengthens a negative result and weakens a positive one.
- **Below the genus the answer stops for a stateable reason.** A phylogenetic random effect within genera costs a tenth of the surviving set; one rank further is impossible on catalogues dereplicated to one genome per species; and on the species pairs that do exist the artefact floor sits sixteenfold above the signal. That is a property of the catalogues, not of the question.
- **Everything is presence and absence.** Copy number, expression and regulation are outside the design, so for claims originally established on read abundance we test whether a gene is carried, not whether it is abundant.
- **Assembly quality is controlled, not eliminated.** No MAG-based comparison can claim the confound is gone. The isolate-collection replication is a partial escape and is reported as partial.

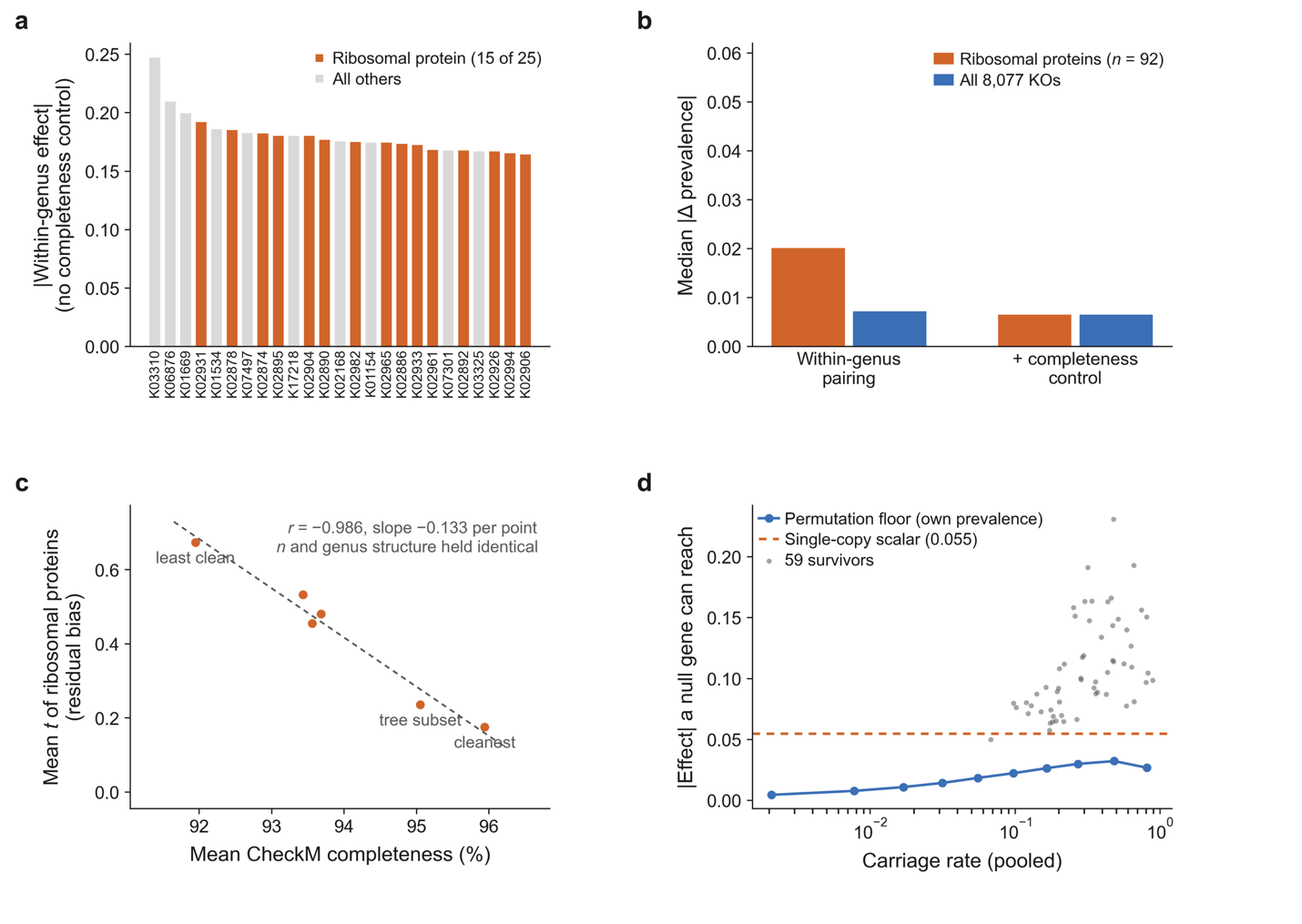

**Extended Data Fig. 1 | The negative control and what its residue is.** **a**, The 25 largest within-genus effects before assembly-quality control, of which the ribosomal proteins are marked; ribosomal proteins cannot truly differ between biomes, so their appearance here shows that within-genus pairing alone does not absorb completeness. **b**, Median |controlled effect| for the 92 ribosomal orthologues against all 8,077 KOs, before and after the quality covariates. **c**, Mean *t* of the ribosomal set against mean CheckM completeness across subsets constructed to hold genome count, genus composition and per-genus biome balance exactly fixed while varying only which genomes are drawn (*r* = −0.986, slope −0.133 per completeness point). The dashed line is ordinary least squares. Because only which genomes are drawn varies, the association cannot be a sample-size effect. **d**, The artefact floor computed two ways: as the 95th percentile of |controlled effect| across the single-copy set, which is the scalar we report and which is drawn as a horizontal rule, and as the 95th percentile of a within-genus permutation null within each of ten bins of pooled carriage rate (2,000 permutations). The permutation floor peaks at mid prevalence, where the sampling null is widest, but its peak is 0.032 against the scalar’s 0.055, because the single-copy set carries the incompleteness bias of panel **c** on top of sampling noise. Points are the 59 survivors at their own carriage rate; all of them clear their own band.

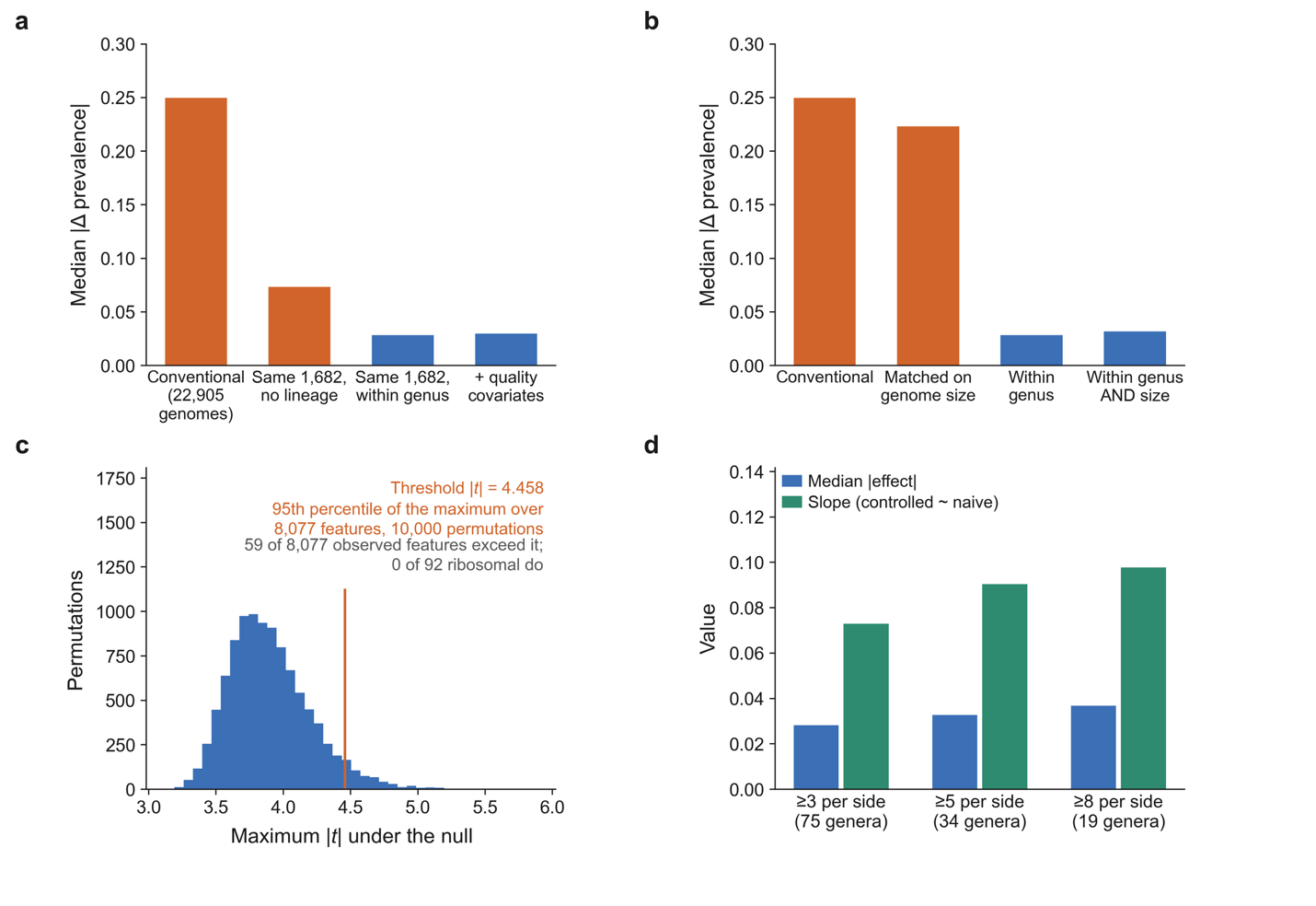

**Extended Data Fig. 2 | Four falsification tests of the collapse.** All panels are the KEGG main specification: 22,905 genomes in the two catalogues, 1,682 of them in the 75 paired genera, 8,077 features, and medians are taken over the 178 features a conventional analysis calls strongly enriched. **a**, Power control: median |Δ| conventionally, then on the same 1,682 paired genomes with no lineage term, then within genus, then adding the quality covariates. If the collapse were lost power the second bar would already show it, and it does not fall to the third. **b**, Genome size: the same quantity with all 22,905 genomes stratified on decile of annotated feature count, which leaves 0.223, and with size decile added on top of genus, which adds nothing to genus alone. Size strata are built on the whole catalogue rather than the paired subset, so that size matching is not confounded with the lineage-overlap selection that defines that subset. **c**, The family-wise null: the distribution of the maximum |*t*| over all 8,077 features across 10,000 within-genus label permutations, with the 95th percentile marked. **d**, Stability across pairing strictness, requiring 3, 5 and 8 genomes per genus per biome, as the median controlled effect and as the slope of controlled on naive; both move upwards as strictness rises, that is, away from zero, so the collapse is not an artefact of admitting weakly paired genera.

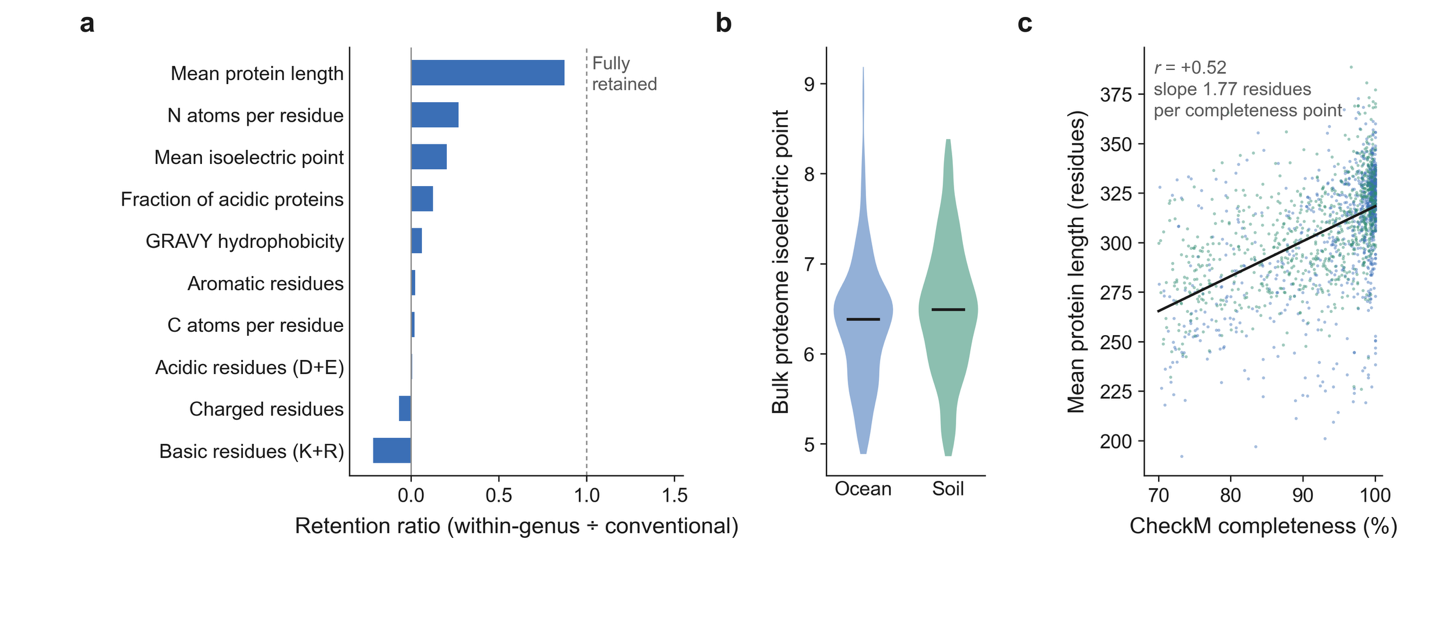

**Extended Data Fig. 3 | Whole-proteome traits.** **a**, Retention ratio (controlled ÷ naive difference) for the ten traits computed over all proteins of each genome, of which nine enter the comparison. Tests are two-sided on the within-genus estimate; the per-trait naive and controlled differences and their *P* values are in the source data for this figure. **b**, Bulk isoelectric point in detail, the trait tested by ref. 46, naive −0.140 against a controlled −0.028 (retention 0.20, *P* = 0.12). **c**, Mean protein length against genome completeness, one point per genome, coloured as in **b** (*r* = +0.52, *P* = 5 × 10⁻¹²⁴, slope 1.77 residues per completeness point, *n* = 1,758). The 4.0-point completeness gap between the catalogues alone predicts a difference of 7.1 residues against the 2.7 observed, which is why this trait is read as assembly fragmentation rather than proteome composition and is excluded from **a**’s median.

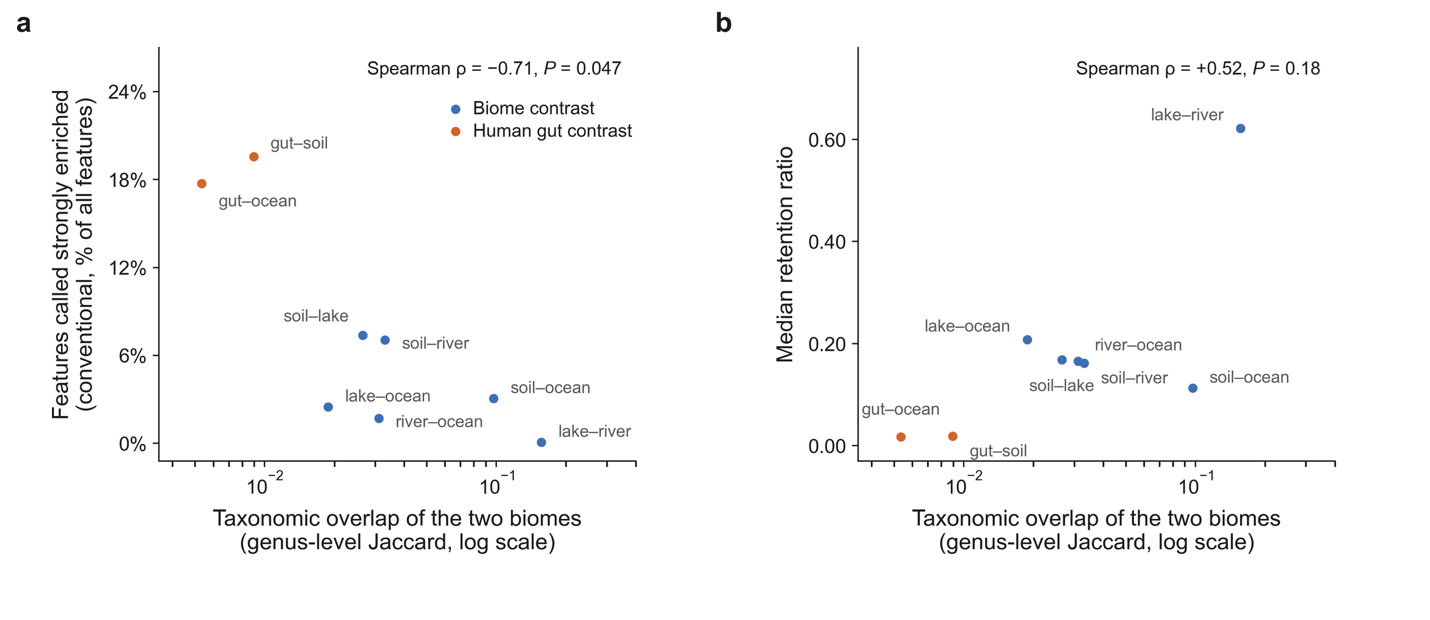

**Extended Data Fig. 4 | Apparent enrichment is ordered by taxonomic overlap.** Fraction of Pfam features called strongly enriched against genus-level Jaccard overlap between the two catalogues, across all eight contrasts (Spearman rho = −0.71, *P* = 0.047, *n* = 8). The corresponding relationship with retention is shown alongside and is not significant (rho = +0.52, *P* = 0.18). Contrasts sharing a catalogue are not independent, so *n* = 8 overstates the degrees of freedom and the relationship is reported as a rule of thumb rather than a calibration.

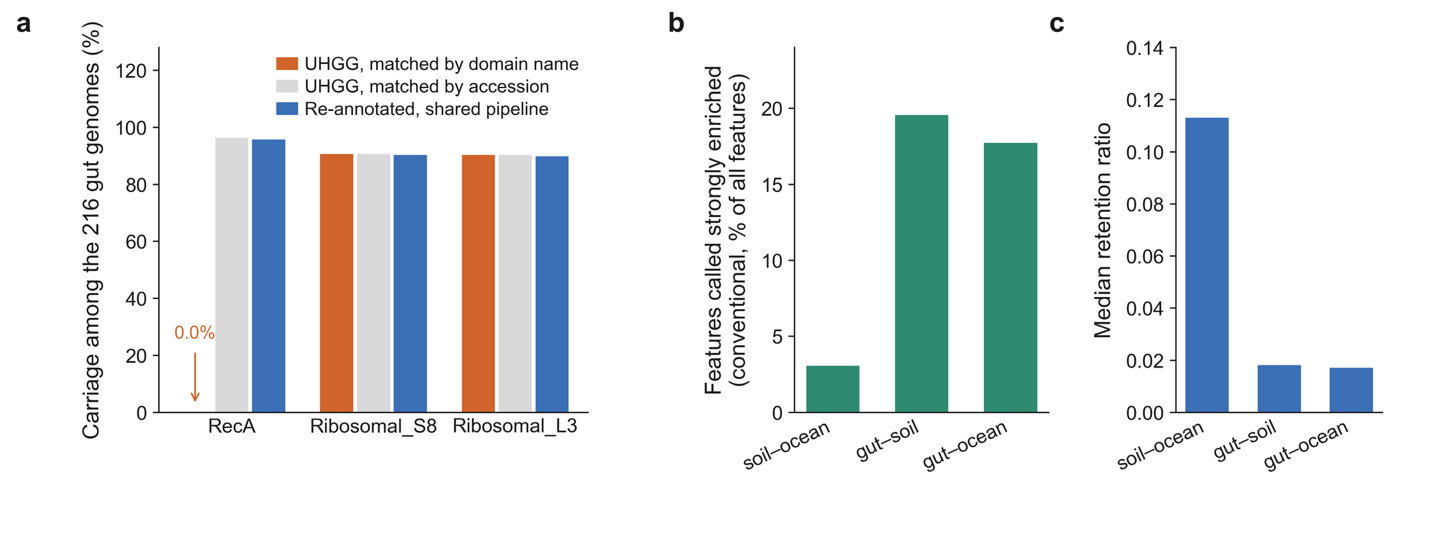

**Extended Data Fig. 5 | The human gut as the limiting case.** **a**, A silent, selective cross-pipeline annotation failure caught by the universal-gene sanity check: under UHGG’s native annotations RecA appears in 0.0% of gut genomes, because UHGG’s profiles are keyed by accession and the current Pfam release names PF00154 RecA_N, so a match on domain name resolves to nothing. Matching on the accession recovers 96.3% and re-annotation through our own pipeline 95.8%. Ribosomal_S8 and Ribosomal_L3 were never renamed and match under either scheme (90.7% and 90.3% natively, 90.3% and 89.8% re-annotated), which is what makes the failure both silent and selective. **b**, Fraction of features called strongly enriched conventionally in the two gut contrasts (13 and 19 paired genera), against the soil–ocean contrast (78 paired genera) as the reference. **c**, Retention after control, 0.02, which both known biases of this catalogue make an upper bound.

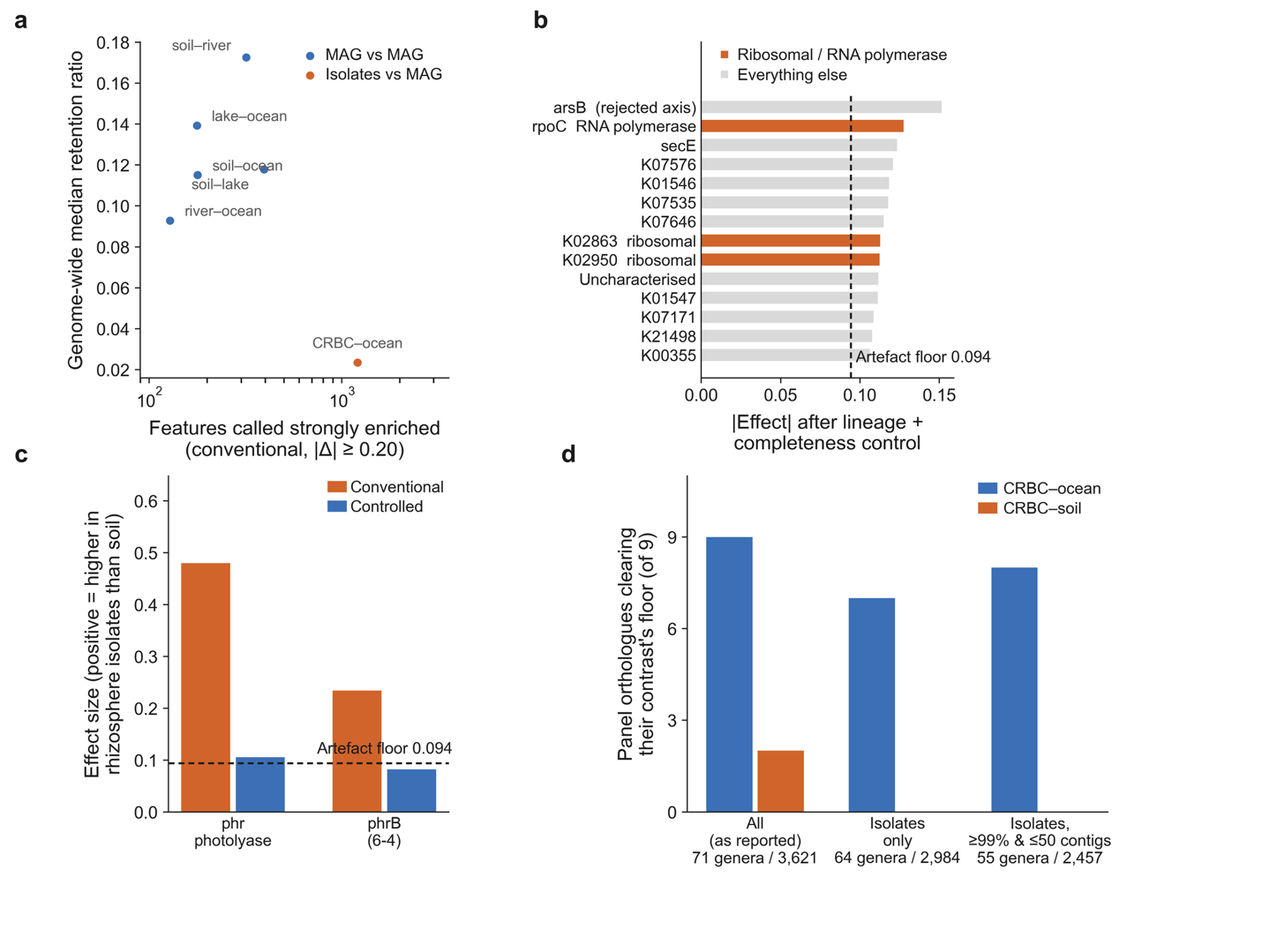

**Extended Data Fig. 6 | The rhizosphere contrast in detail.** **a**, Why genome-wide median retention is not comparable across contrasts of differing compositional distance: 733 to 1,258 features reach naive |Δ| ≥ 0.20 in the CRBC contrasts against 178 for soil, ocean, and the greater the compositional distance the lower the genome-wide retention. **b**, The residual technical tail in the CRBC–soil comparison, in which no biome contrast exists, with that contrast’s artefact floor of 0.094 (95% CI 0.078 to 0.112) marked. The genome-wide median controlled effect is pressed to 0.007, but the fourteen largest all clear the floor, at |Δ| = 0.106 to 0.151; three are RNA polymerase or ribosomal, the largest is arsB, an axis this paper has already rejected, and the rest are uncharacterised. Isolate-versus-MAG assembly quality is therefore not fully absorbed. **c**, The photolyase observation against the floor that decides it. phrB falls below the floor (0.082 against 0.094) and phr sits just above it (0.105); neither clears the floor’s upper confidence bound of 0.112, which is why the observation is withdrawn. Effects and floor are drawn from the same quality-subset table so that both are on one specification. **d**, Nested quality subsets: isolate assemblies only (64 genera, 2,984 genomes) and isolates at completeness ≥ 99% with ≤ 50 contigs (55 genera, 2,457 genomes) give 9, 7 and 8 of the nine panel orthologues clearing their contrast’s floor in CRBC–ocean, against 2, 0 and 0 in CRBC–soil. Genome-wide retention is 0.017 to 0.025 throughout. Tightening quality raises the floor rather than lowering it, because the subsets are smaller and the estimates noisier.

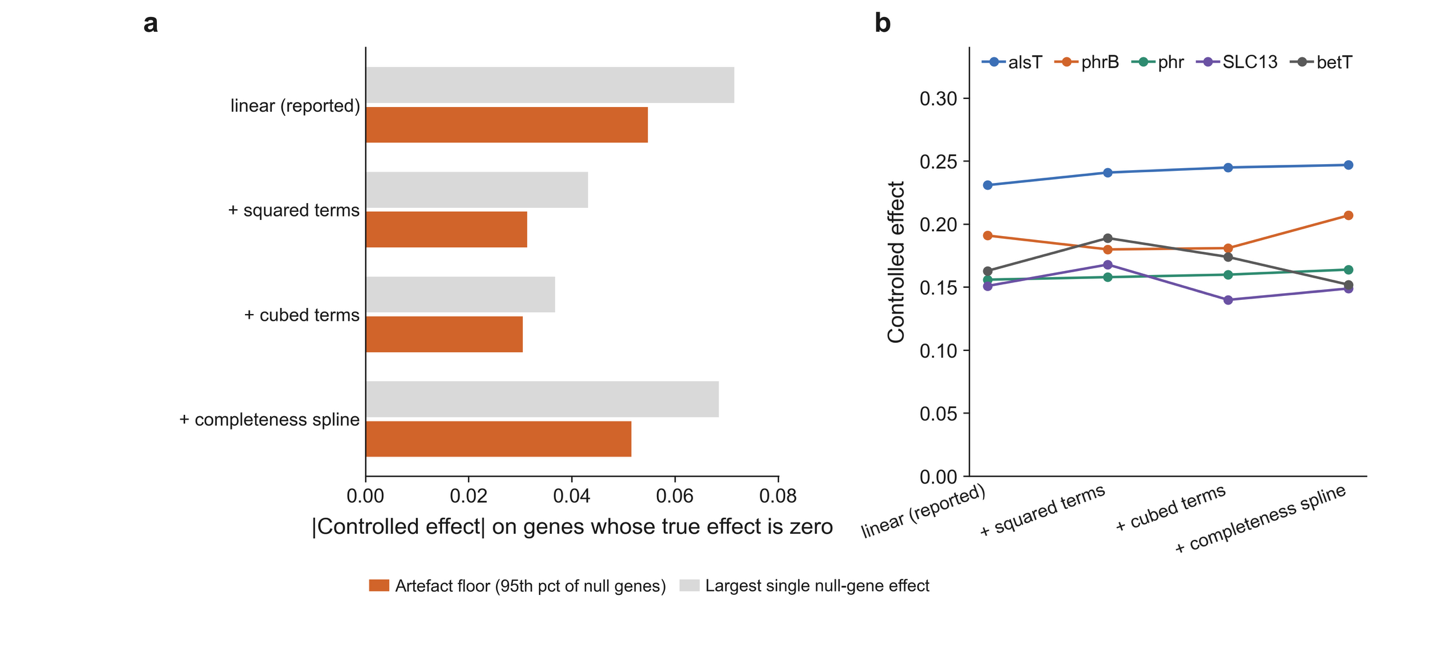

**Extended Data Fig. 7 | Non-linear quality covariates tighten the negative control without touching the signal.** Artefact floor, ribosomal maximum and the five largest surviving effects under the linear specification reported throughout and under specifications adding squared, cubed and piecewise-linear terms in completeness and feature count (1,682 genomes, 75 genera in every row). Squared terms move the floor from 0.055 to 0.031 while every surviving effect holds or grows, which identifies the residue on the negative control as assembly non-linearity rather than biology. The linear specification is retained because it leaves the floor roughly twice as high and is therefore the conservative choice.

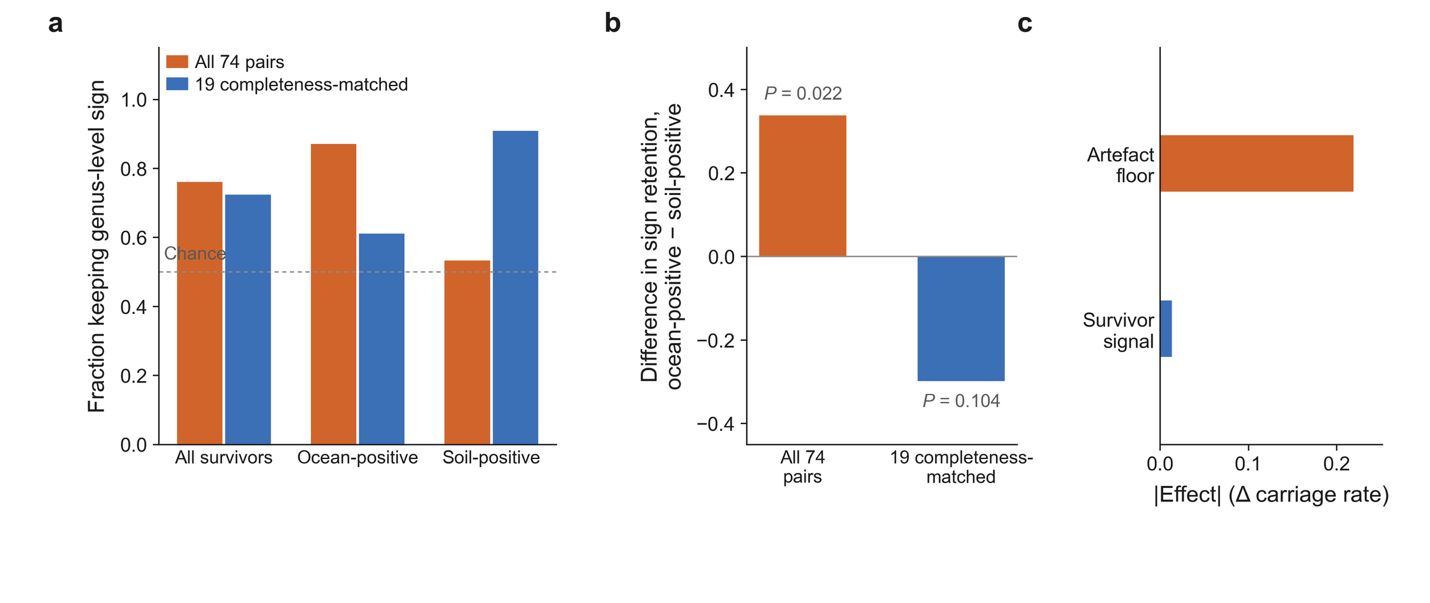

**Extended Data Fig. 8 | The species-level directional result is a completeness artefact.** The evidence behind a withdrawal; nothing here is offered as a positive finding. **a**, Fraction of the 59 genus-level survivors keeping their sign in the paired within-species comparison, for all 74 species pairs and for the 19 pairs matched on completeness to within one percentage point, split by the side the genus-level effect favoured. Features whose within-species effect is exactly zero carry no directional information and are excluded (13 of 59 in the full set, 30 of 59 in the matched subset); the per-bar numerators and denominators are in the source data for this figure. The dashed rule is the matched-random expectation. **b**, The difference between the ocean-positive and soil-positive retention rates, with permutation *P* values from 10,000 label swaps. In the full set the asymmetry is +0.34 (*P* = 0.022), which is what a directional completeness gap of +3.7 percentage points would produce; in the completeness-matched pairs it is −0.30 (*P* = 0.10), that is, absent. **c**, The artefact floor at this level, 0.219, against the median |within-species effect| of the 59 survivors, 0.0135, a factor of 16. Power is not the constraint: at a median standard error of 0.013 a genus-level effect of 0.098 carried intact would have been detected.
